# Development of PTSD-like behavior in adult mice after observing an acute traumatic event

**DOI:** 10.1101/249870

**Authors:** Ray X. Lee, Greg J. Stephens, Bernd Kuhn

## Abstract

In human post-traumatic stress disorder (PTSD), a major psychiatry challenge is how diverse stress reactions emerge after a protracted symptom-free period. Here, we study the behavioral development in mice isolated after observing an aggressive encounter inflicted upon their pair-housed partners and compared the results with those in multiple control paradigms. Compared with mice plainly isolated, mice isolated following the acute witnessing social stress gradually developed a wide range of long-term differences of their physiological conditions, spontaneous behaviors, and social interactions, including paradoxical results if interpreted in traditional ways. To address this developmental diversity, we applied fine-scale behavioral analysis to standard behavioral tests and showed that the seemingly sudden emergent behavioral differences developed gradually. Mice showed different developmental patterns in different zones of a behavior testing apparatus. However, the results of the fine-scale analysis together with state-space behavioral characterization allow a consistent interpretation of the seemingly conflicting observations among multiple tests. Interestingly, these behavioral differences were not observed if the aggressive encounter happened to a stranger mouse. Additionally, traumatized mice showed rebound responses to their partners after the long separation. In contrast, mice pair-housed with their attacked partners after the aggressive encounters still showed a difference in social interactions, while a difference in spontaneous behaviors did not occur. Accordingly, we propose that social relationship is the single common factor underlying the otherwise independent development of behavioral differences in this mouse paradigm and that the gained insights could have parallels in human PTSD development.

## Introduction

Stress reactions can emerge long after the triggering event. Stress incubation describes the time interval following an aversive event during which stress reactions emerge or increase. Since the phenomenon of anxiety and fear incubation was first formulated and summarized (Diven, 1937), stress incubation has been assumed to be spontaneous in the sense that the development of behavioral changes is highly determined by internal rather than external causes (McAllister DE, 1967). The phenomenon of stress incubation has received serious clinical and research attention in human psychiatry, especially characterized in the post-traumatic stress disorder (PTSD), one of the most prevalent mental health disorders (DSM-5, 2013; DSM-III, 1980). Despite such prevalence, PTSD remains far from understood and controversial (McFarlane, 2010). The debates have even questioned the existence of PTSD (McHugh and Treisman, 2007; Walton et al., 2017), due to its diversity, inconsistency, and delayed onset of symptoms even after a protracted symptom-free period (Andrews et al., 2007; Pai et al., 2017).

To identify the psychopathological developments during stress incubation, it is beneficial to use an experimental assay with purely psychosocial manipulations on controlled subjective experiences and a homogeneous genetic background. Laboratory rodents have been used to study stress behavior and the pharmacology of stress (Calhoon and Tye, 2015; Cryan and Holmes, 2005; Kaouane et al., 2012; Tovote et al., 2015). A delay period before showing substantial stress reactions, suggesting stress incubation, was reported in the context of rodent models simulating human PTSD (Davis, 1989; Pamplona et al., 2011; Sillivan et al., 2017; Tsuda et al., 2015; Warren et al., 2013). The wide variety of aversive stimuli in these models range from acute physical stress (Balogh et al., 2002; Philbert et al., 2011) to prolonged witnessing of social defeat (Sial et al., 2016; Warren et al., 2013). Compared with mice experiencing direct social attacks, mice observing social attacks showed a more obvious phenomenon of stress incubation (Warren et al., 2013). This observation emphasizes the significance of emotional and cognitive processes, but not the direct impact of physical stress, in stress incubation (Hayes et al., 2012). Still, many major questions remain unanswered: Why do some symptoms attenuate while the others incubate? (Bryant et al., 2017, 2013) Why does a detailed difference in subjective experiences lead to dramatic variation in stress developments? (Adams and Boscarino, 2006) Why are symptoms usually treated partially within varied recovery contexts? (Harvey, 1996) Why does a treatment that rescues one’s symptom even worsen the same symptom of another? (Scherer et al., 2017) Is stress incubation different from the formation, storage, and retrieval of fear memory? (Poulos et al., 2014)

To address such questions, we need a better understanding of the behavioral correlates in stress incubation. Here, we combined behavioral scenarios, analytic methods, and psychological hypotheses to study the development of stress incubation by psychobehavioral experiments in mice. We systematically, quantitatively, and longitudinally examined multiple physiological conditions (body mass, corticosterone level, brain connectome, etc.), spontaneous behaviors (light-dark box, open field, locomotion, etc.), and social interactions (female strangers, male strangers, pair-housed partners, etc.) of mice after observing acute social stress happening to their familiar partners (Figure 1A), with seven relevant control paradigms (Figure 1, B to H). Along the investigation, we introduced methods of fine-scale behavioral analysis and state-space behavioral characterization to overcome paradoxical and inconclusive results commonly observed in the traditional analyses of standard behavioral tests when the speculated emotions underlying behaviors are subtle and complex (Ramos, 2008). Lastly, we discussed a potential conceptual model of psychological framework based on our tests and observations in mouse stress incubation, which might provide insights on prevention, detection, and treatments of human PTSD development.

**Figure 1.**
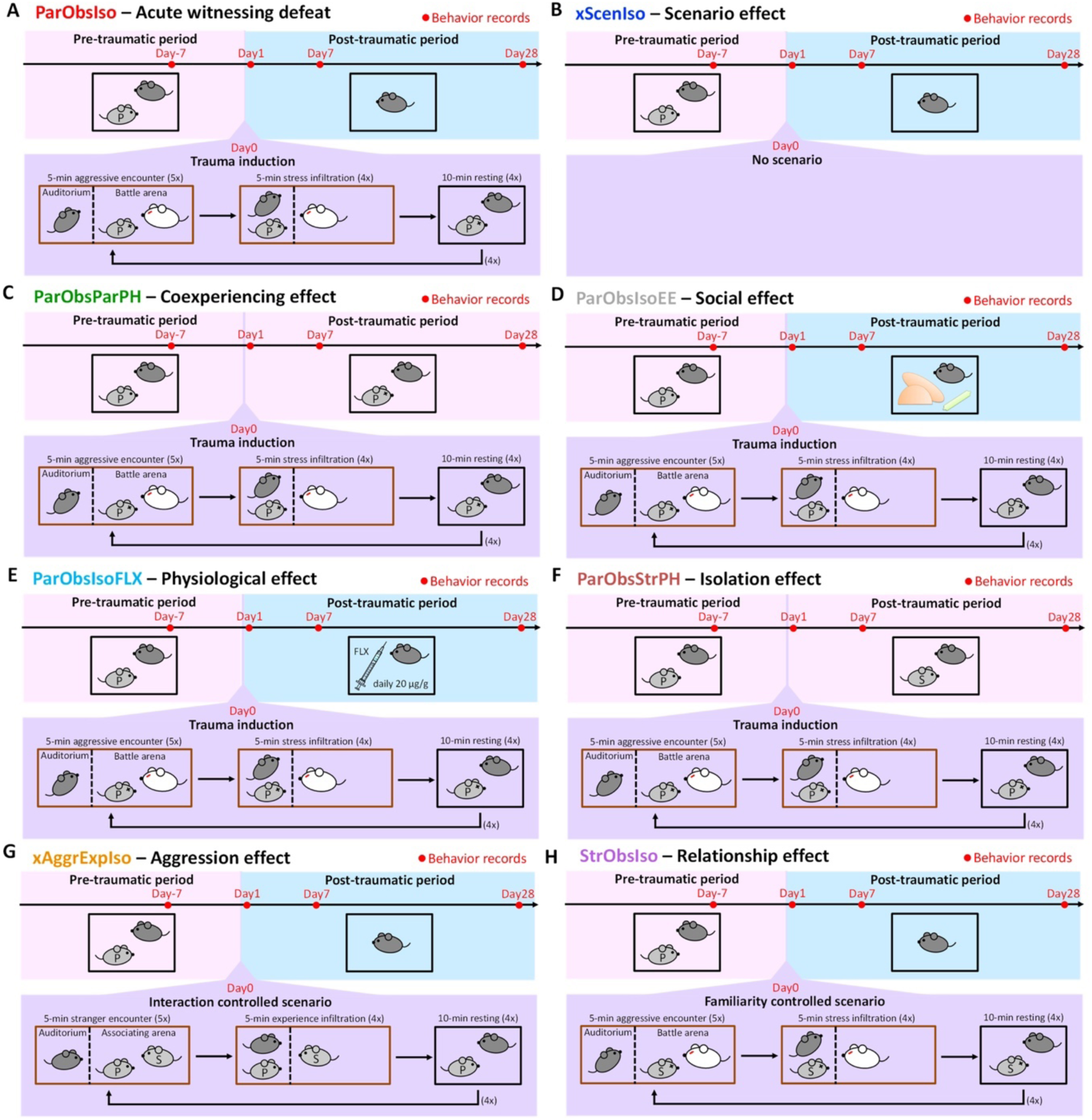
Paradigm inducing PTSD-like behavior in mice and seven control paradigms. **(A)** Paradigm with acute psychosocial trauma induction in mice. Focal mouse [dark gray; Partner-Observing-Isolated (ParObsIso) mouse], partner mouse (light gray P), aggressor mouse (white), focal mouse’s homecage (black), aggressor’s homecage (brown), and wire-meshed divider (dashed line). **(B)** No-Scenario-Isolated (xScenIso) mice were separated without trauma induction and identified the scenario effect in the behavioral paradigm. **(C)** Partner-Observing-Partner-Pair-Housed (ParObsParPH) mice were pair-housed with their partners after trauma induction and identified the social transfer effect of co-experiencing trauma in the behavioral paradigm. **(D)** Partner-Observing-Isolated-Environment-Enriched (ParObsIsoEE) mice were provided with toys after trauma induction and identified the social rescue effect in the behavioral paradigm. **(E)** Partner-Observing-Isolated-Fluoxetine-treated (ParObsIsoFLX) mice were treated with fluoxetine after trauma induction and identified the pharmacological rescue effect in the behavioral paradigm. **(F)** Partner-Observing-Stranger-Pair-Housed (ParObsStrPH) mice were pair-housed with strangers after trauma induction and identified the isolation effect in the behavioral paradigm. **(G)** Non-Aggressor-Exposed-Isolated (xAggrExpIso) mice had experience of social interactions without witnessing stress from strangers and identified the aggression effect in the behavioral paradigm. **(H)** Stranger-Observing-Isolated (StrObsIso) mice had witnessing experience of trauma that happened to strangers, rather than to their pair-housed partners, and identified the relationship effect in the behavioral paradigm.

## Results

### Long-term effects emerged after acute trauma induction

To identify post-traumatic behavioral development with minimal physical impact and minimal peri-traumatic effects, we developed the following assay of an acute witnessing trauma (Figure 1A): Partnership between the male focal mouse and its male partner was established by housing them together for 3 weeks (Day-21–Day0). During this pair-housing, the mice slept together in a single nest which they built and no aggressive interaction (attacks, pursuits, and over-allogrooming) was observed. On Day0 (trauma induction), the focal mouse observed its partner being attacked by 5 different aggressor mice in succession (aggressive encounters) and stayed together with the attacked partner between each aggressive encounter (stress infiltration and resting). Importantly, the focal mouse only experienced the traumatic event by witnessing the attacks and by social communication and olfactory cues, but not through any direct physical threat, such as attack bites and pursuits from either the aggressors or its partner. After the last aggressive encounter, the focal observer mouse [Partner-Observing-Isolated (ParObsIso) mouse] was socially isolated for 4 weeks (Day0–Day28). Behavior was tested on Days -7, 1, 7, and 28.

To differentiate behavioral consequences from the trauma induction and the effects of isolation, adaptation to the tests, and aging, we first compared ParObsIso mice with a control group of mice isolated from their partners on Day0 without trauma induction [No-Scenario-Isolated (xScenIso) mice; Figure 1B]. We found that ParObsIso mice (n = 47) built nests with significantly higher walls than those constructed by xScenIso mice (n = 47) after isolation (Figure 2A). ParObsIso mice also increased their body mass in the late phase of the study (Figure 2B). Both observations indicate a long-term effect of the trauma induction.

**Figure 2.**
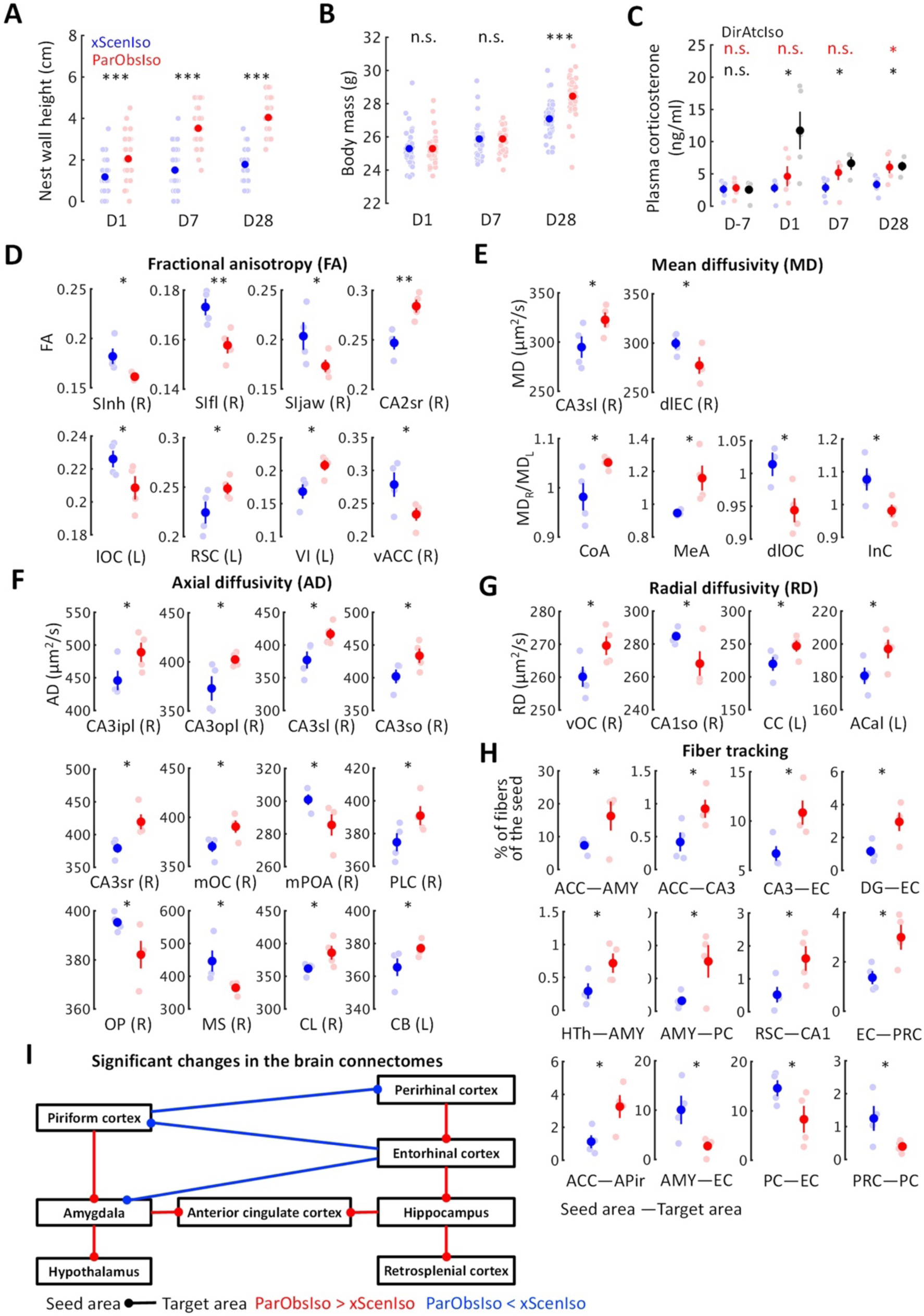
Long-term and delayed effects on multiple behaviors and physical conditions. **(A)** Nest wall heights show long-lasting significant differences after trauma induction. **(B)** Body mass shows a significant increase 28 days after trauma induction. **(C)** Baseline plasma corticosterone level increased after trauma induction for both ParObsIso mice and their partners, the DirAtcIso mice. **(D)** Fractional anisotropy (FA) of DTI-based water diffusivity suggests the changes of average microstructural integrity in multiple areas of the cerebral cortex. lOC, lateral orbital cortex; SInh, non-homunculus region of the primary sensory cortex; SIfl, forelimb region of the primary sensory cortex; SIjaw, jaw region of the primary sensory cortex; vACC, ventral region of the anterior cingulate cortex; CA2sr, stratum radiatum of the hippocampal cornu ammonis (CA) 2 area; RSC, the retrosplenial cortex; VI, the primary visual cortex; (R), the right area; (L), the left area. **(E)** DTI-based mean water diffusivity (MD) suggests the changes of membrane density in multiple areas of the entorhinal cortex-hippocampus system and the straitening of structural hemispheric specializations in the amygdala-insular cortex system. dlOC, dorsolateral orbital cortex; InC, the insular cortex; CoA, the cortical amygdalar nucleus; MeA, medial amygdalar nucleus; CA3sl, stratum lucidum of the hippocampal CA3 area; dlEC, dorsolateral entorhinal cortex. **(F)** DTI-based axial water diffusivity (AD) suggests the changes of neurite organization in multiple areas of the cerebral cortex and white matter mainly in the right hemisphere. OP, olfactory peduncle; PLC, prelimbic cortex; mOC, medial orbital cortex; CL, claustrum; MS, medial septal complex; mPOA, medial preoptic area; CB, cingulum bundle; CA3sr, stratum radiatum of the hippocampal CA3 area; CA3sl, stratum lucidum of the hippocampal CA3 area;CA3so, stratum oriens of the hippocampal CA3 area; CA3ipl, inner pyramidal layer of the hippocampal CA3 area; CA3opl, outer pyramidal layer of the hippocampal CA3 area. **(G)** DTI-based radial water diffusivity (RD) suggests the changes of myelination in multiple areas of the cerebral cortex in the right hemisphere and the white matter in the left hemisphere. vOC, ventral orbital cortex; ACal, anterior limb of the anterior commissure; CC, corpus callosum; CA1so, stratum oriens of the hippocampal CA1 area. **(H)** DTI-based network-wise fiber tracking reveals specific chronic changes of structural connectivity in the brain. AMY, the amygdala; HTh, the hypothalamus; DG, the hippocampal dentate gyrus; PC, the piriform cortex; PRC, the perirhinal cortex; APir, the amygdalopiriform transition area. Note that the brain regions were in the right brain hemisphere. **(I)** Trauma-induced structural changes of the underlying brain connectome revealed a network enhancement centered at the anterior cingulate cortex. Error bars indicate standard errors of the means; n.s., p≥0.05; *, 0.01≤p<0.05; ***, p<0.001; two-sample Student’s t-test.

To further explore potential physiological changes related to stress underlying the paradigm, we examined corticosterone concentrations in blood plasma. Compared with xScenIso mice, ParObsIso mice showed higher baseline plasma corticosterone level (CORT) after trauma, which reached statistical significance on Day28 (Figure 2C). In this experiment, we also compared CORT of ParObsIso mice with that of their partners, the attacked mice isolated after trauma [Directly-attacked-isolated (DirAtcIso) mice; n=5, note that 2 out of 5 mice died on Days 4 and 5, respectively, without obvious physical trauma; interestingly, such losses were not observed in the directly attacked mice which were subsequently group-housed]. The tendency of higher CORT was also observed in DirAtcIso mice which had a more obvious CORT increment during the early phase (Figure 2C), indicating that the observed differences of nest wall height and body mass may be phenotypes of stress.

To obtain indication of microstructural changes in the full brain correlated with these physiological and behavioral differences, we used diffusion tensor imaging (DTI) of brains collected on Day28 to analyze brain-wide microstructural differences (Figure 2, D to G; Supplemental Figures 1 to 4): Rather than a structural change in the hypothalamus which modulates CORT (DeMorrow et al., 2018), significant differences mainly occurred in both neocortex and hippocampus. For ParObsIso mice, compared with xScenIso mice, their DTI-based fractional anisotropy (FA; Figure 2D) was lower in anterior cerebral cortical areas including the primary somatosensory cortex, anterior cingulate cortex, and orbital cortex, but higher in posterior cerebral cortical areas including the retrosplenial cortex and the primary visual cortex. It was also higher in the hippocampus. Different measures of DTI-based water diffusivities [mean diffusivity (MD), axial diffusivity (AD), and radial diffusivity (RD); Figure 2, E to G] were generally higher in the cerebral cortex and hippocampus of ParObsIso than in xScenIso mice. Interestingly, we observed an obvious asymmetry of the brain areas: Gray matters including the cerebral cortex and hippocampus showed differences in the right brain hemisphere, while white matters including the corpus callosum, anterior commissure, and cingulum bundle showed differences in the left brain hemisphere. The asymmetry of diffusivity between the right and left brain hemispheres were most significant in the amygdala (right hemisphere shows higher diffusivity) and areas of the lateral cerebral cortical subnetwork including dorsolateral orbital cortex and insular cortex (left hemisphere shows higher diffusivity) for ParObsIso mice.

For long-range connections, the brains of ParObsIso mice show a significant increase of structural connections between the right hippocampus (CA3 and dentate gyrus) and right dorsolateral entorhinal cortex (Figure 2H) compared to the brain of xScenIso mice. While the entorhinal cortex is the main interface between the hippocampus and neocortex, we further tracked its long-range connections. The right dorsolateral entorhinal cortex showed significantly increased connections with the right perirhinal cortex, but had significantly decreased connections with the right piriform cortex (Figure 2H). Between the piriform and perirhinal cortices, we found a decrease of their connections (Figure 2H). Through brain-wide tracking, we revealed trauma-induced changes in the “perirhinal cortex–entorhinal cortex– hippocampus” system, the “retrosplenial cortex–hippocampus” system, the “piriform cortex– amyglala” system, and the “amyglala–hypothalamus” system. All of them centered at the anterior cingulate cortex (ACC; Figure 2I), suggesting that the emergence of long-term behavioral changes after trauma induction arise from network changes in this high-level cognitive and emotional center.

### Fine-scale behavioral analysis revealed gradual stress development

Spontaneous behaviors in a steady environment has been taken as a representation of internal states such as emotion (Archer, 1973). Behavioral tests for rodent anxiety-like reactions were designed to evaluate their stress reaction against their willingness to explore (e.g. light-dark box and elevated plus-maze) or their activity under conditions with a gradient of uncertainty (e.g. open field). We first examined spontaneous behaviors in the light-dark box test (Figure 3A; n = 8 mice for each group), where the stressor was a natural aversion to brightly lit areas (Kumar et al., 2013). While the time spent in the light area did not differ significantly between xScenIso and ParObsIso mice on Day1, ParObsIso mice surprisingly spent more time in the light area than xScenIso mice on Days 7 and 28 (Figure 3A, left panels). This result raised the following questions: (i) Did this behavioral difference start to develop immediately after the traumatic event or only after a delay? (ii) What comparative emotionality does this behavioral difference indicate?

**Figure 3.**
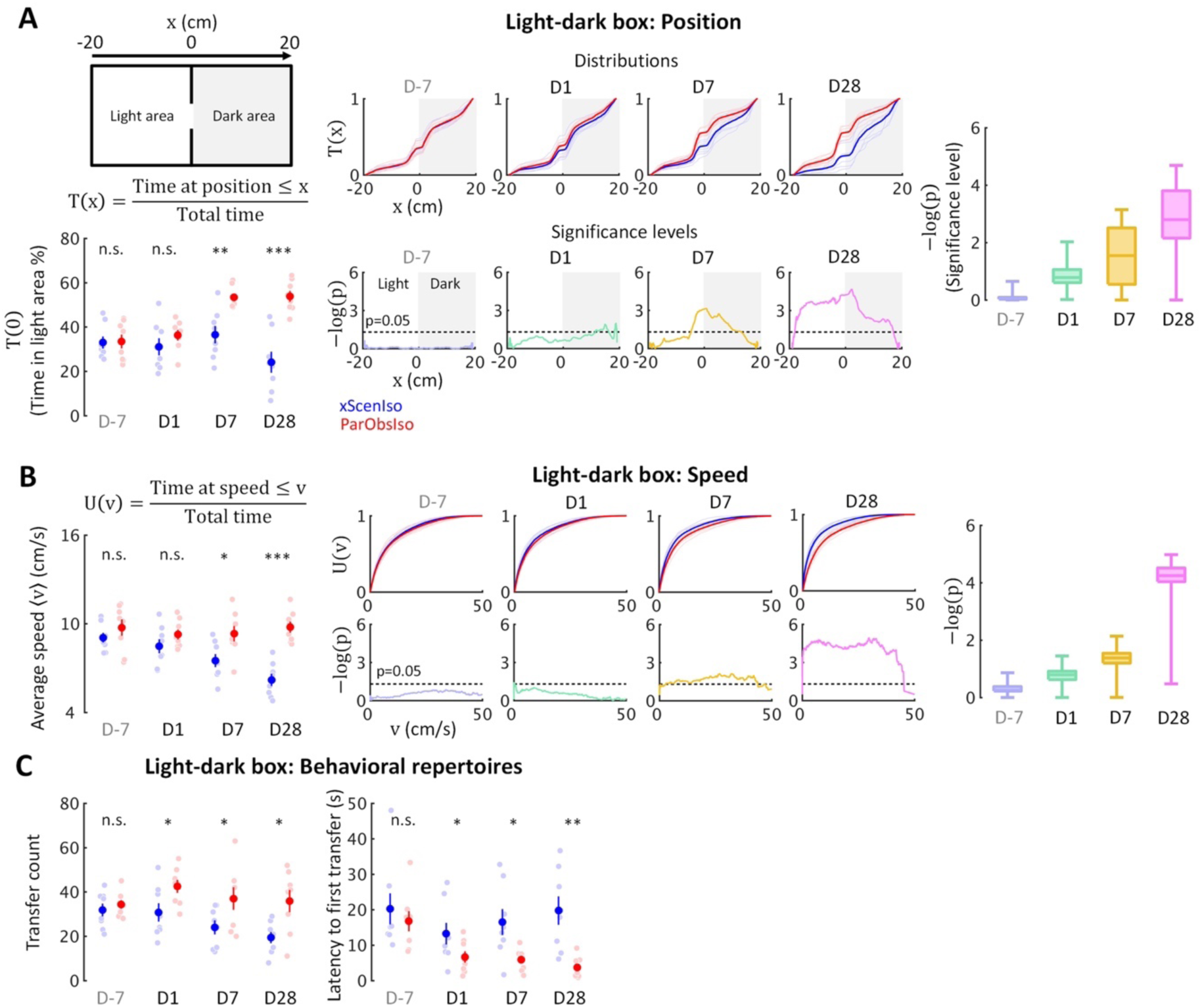
Fine-scale behavioral analysis in light-dark box test detects the gradually developing process of the behavioral difference. **(A)** Light-dark box test quantified through the cumulative position probability T(x) along the light-dark axis (left; total time = 300 s). On average, ParObsIso mice (red) spent more time in the light area than xScenIso mice (blue) during the late post-traumatic period [T(0), bottom-left]. Spatially fine-scale behavioral analysis reveals significant differences between ParObsIso and xScenIso populations already in the early post-traumatic period (middle). For each position, we compute the mean T(x) across the xScenIso and ParObsIso populations and compute statistical significance through a two-population Student’s t-test. These differences gradually increased, as evidenced by significance distributions collapsed across all positions (right; box plots show the minima, lower quartiles, medians, upper quartiles, and maxima). **(B)** We similarly quantified speed using the fine-scale cumulative distribution U(v)of having speed ≤v and we show the statistical analysis of population differences in U(v). Cumulative distribution functions of locomotion speed (U(v) of having speed ≤v) and corresponding significance distributions provide an additional independent behavioral index that showed a gradually increasing differences of higher speed in ParObsIso mice. **(C)** Higher transfer counts and shorter latency to the first transfers in ParObsIso mice suggest their higher activity and exploratory motivation, respectively.

To answer (i), we examined the positions of the mice in the light-dark box on a fine-scale. Based on spatial symmetry, we analyzed T(x), the cumulative probability distribution of time that the mouse spent at positions along the axis of the light-dark box (Figure 4A, top-right scheme and the equation, and middle panels for the results). We calculated significance levels [presented as -log of p-values, -log(p)] of T(x) between ParObsIso and xScenIso populations by computing position-dependent population means and applying a two-tailed, two-sample Student’s t-test (Figure 3A, bottom-middle panels). Already on Day1, ParObsIso mice showed differences in their spatial distribution, as they spent less time than xScenIso mice at the far end of the dark area. This tendency increased with time: On Day 7, ParObsIso mice spent more time in the light area close to the door compared to xScenIso, and then on Day 28 additionally on the far side of the light area. Collapsed -log(p) distributions reveal the overall gradual increase in spatial preference differences (Figure 3A, right panel). Additionally, ParObsIso mice maintained a higher locomotor speed compared to the gradually decreasing speed of xScenIso mice (Figure 3B) and showed more transfers between the boxes and shorter latencies until their first transfers from Day 1 (Figure 3C). These results indicate an onset and gradual increase of behavioral differences immediately following the trauma.

**Figure 4.**
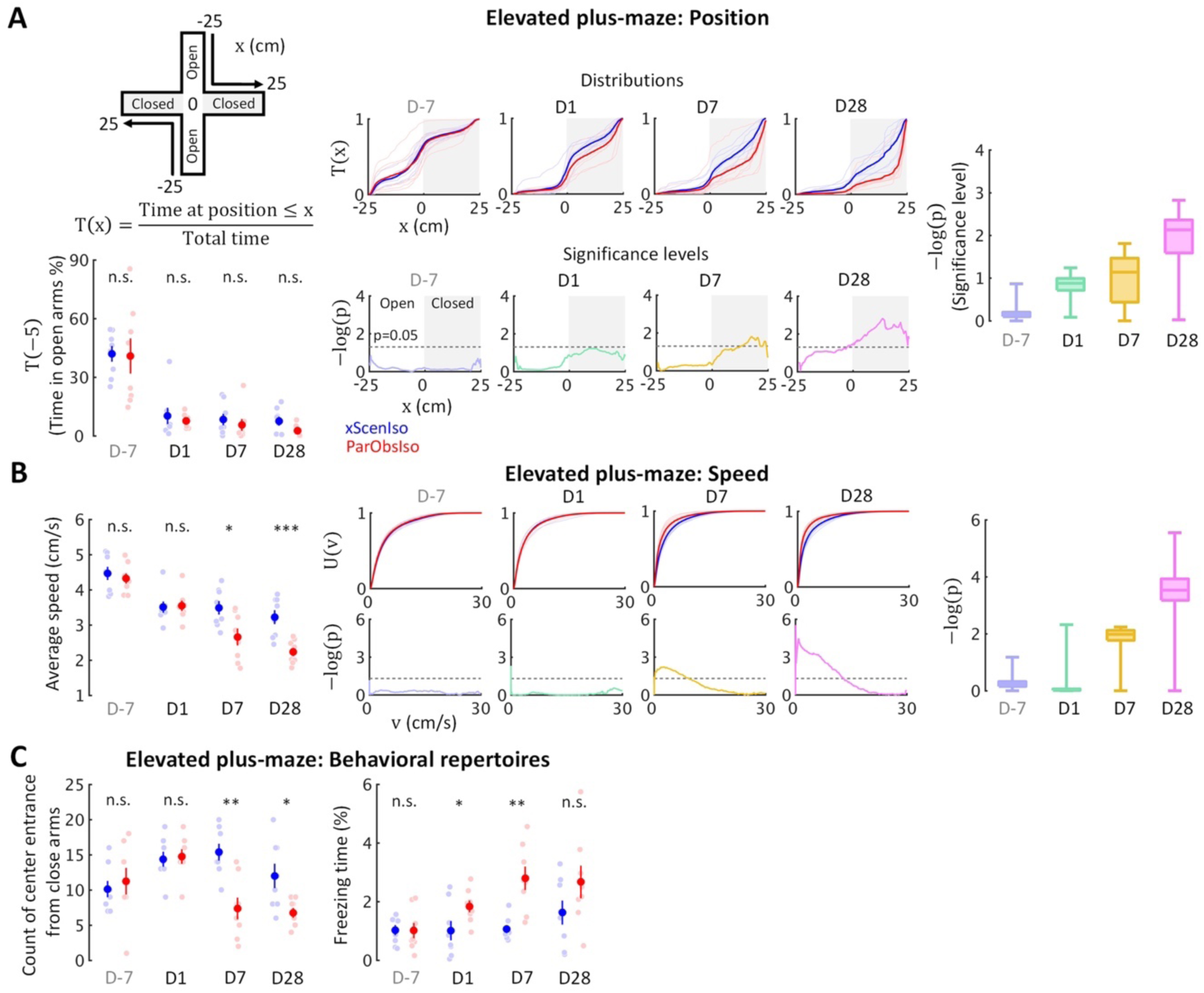
Behavioral testing in the elevated plus-maze demonstrates that stress incubation of anxiety caused the observed differences. **(A)** ParObsIso and xScenIso mice did not differ significantly in the time they spent in opened arms (total time = 300 s); however, spatial distributions show differences in preferred location between ParObsIso and xScenIso mice in the closed arms, which increased with time. **(B)** Cumulative distribution functions of locomotion speed and corresponding significance distributions show a gradually increasing differences of lower speeds in ParObsIso mice. **(C)** Less exploration from close arms to platform center and longer freezing time in ParObsIso mice suggest their stronger stress reactions.

Regarding (ii), we additionally examined their behaviors in the elevated plus-maze test (Figure 4A; n = 8 mice for each group). In this test, stressors included fear of falling and exposure. After first exposure on Day-7 and separation, mice spent only a fraction of the time in the open arms of the maze, but with no significant difference between xScenIso and ParObsIso mice (Figure 4A, left panels). However, ParObsIso mice spent increasingly more time in the far end of the closed arms (Figure 4A, middle panels) and moved more slowly in the elevated plus-maze after trauma induction (Figure 4B) with longer periods of freezing and fewer entries to the central platform from the closed arms (Figure 4C). Although the gradually increasing differences between xScenIso and ParObsIso mice (Figure 4, A and B) was consistent, the opposite tendency of reactions in light-dark box and elevated plus-maze tests was seemingly paradoxical.

Seemingly conflicted results are regularly observed in traditional analyses of standard behavioral tests as the emotions underlying behaviors may be subtle and complex (Ramos, 2008; Ramos et al., 2008). To investigate possible psychological interpretation of the opposite reactions, we compared the behaviors of xScenIso and ParObsIso mice in the tests with the tested behaviors of mice injected with caffeine, which induces anxiety somatically (DSM-5, 2013), and mice after experiencing brief shocks, which induces anxiety cognitively (Bolton and Robinson, 2017), under the otherwise same experimental conditions and procedures. Reactions of cognitive anxiety are expected to be opposite of those shown in somatic anxiety (Cloninger, 1988). In addition, the behavioral characteristics of caffeine-injected mice and foot-shocked mice in standard tests remain controversial (Borsini et al., 2002; Gulick and Gould, 2009; Suarez and Gallup, 1981; Wu et al., 2017), as what we also found when applying traditional analyses (Figure 5, A and B). To capture behavioral characteristics that may be overlooked in a subjective, low-dimensional representation, we first evaluated the local likelihood of a given behavioral state (described by position, instantaneous locomotor speed, instantaneous locomotor acceleration, and instantaneous velocity along the stressor-to-stressor-free axis) to be recorded from caffeine-injected, foot-shocked, or non-treated mice. Behavioral characteristics that are consistent within a group but varied across groups were therefore quantitatively identified. By referring local likelihoods distributing across entire behavioral state space (Supplemental Videos 1–6), we calculated the global likelihood of behaviors of xScenIso and ParObsIso mice in a test to be caffeine-injected-like, foot-shocked-like, or non-treated-like. While xScenIso mice kept showing non-treated-like behaviors in both tests although their behaviors in classical analyses changed, ParObsIso mice developed caffeine-injected-like behaviors in the light-dark box test and developed foot-shocked-like behaviors in the elevated plus-maze test after trauma induction (Figure 5, C and D). The results suggest that ParObsIso mice display more behaviors of chronic somatic anxiety when tested in the light-dark box test and more behaviors of chronic cognitive anxiety when tested in the elevated plus-maze test.

**Figure 5.**
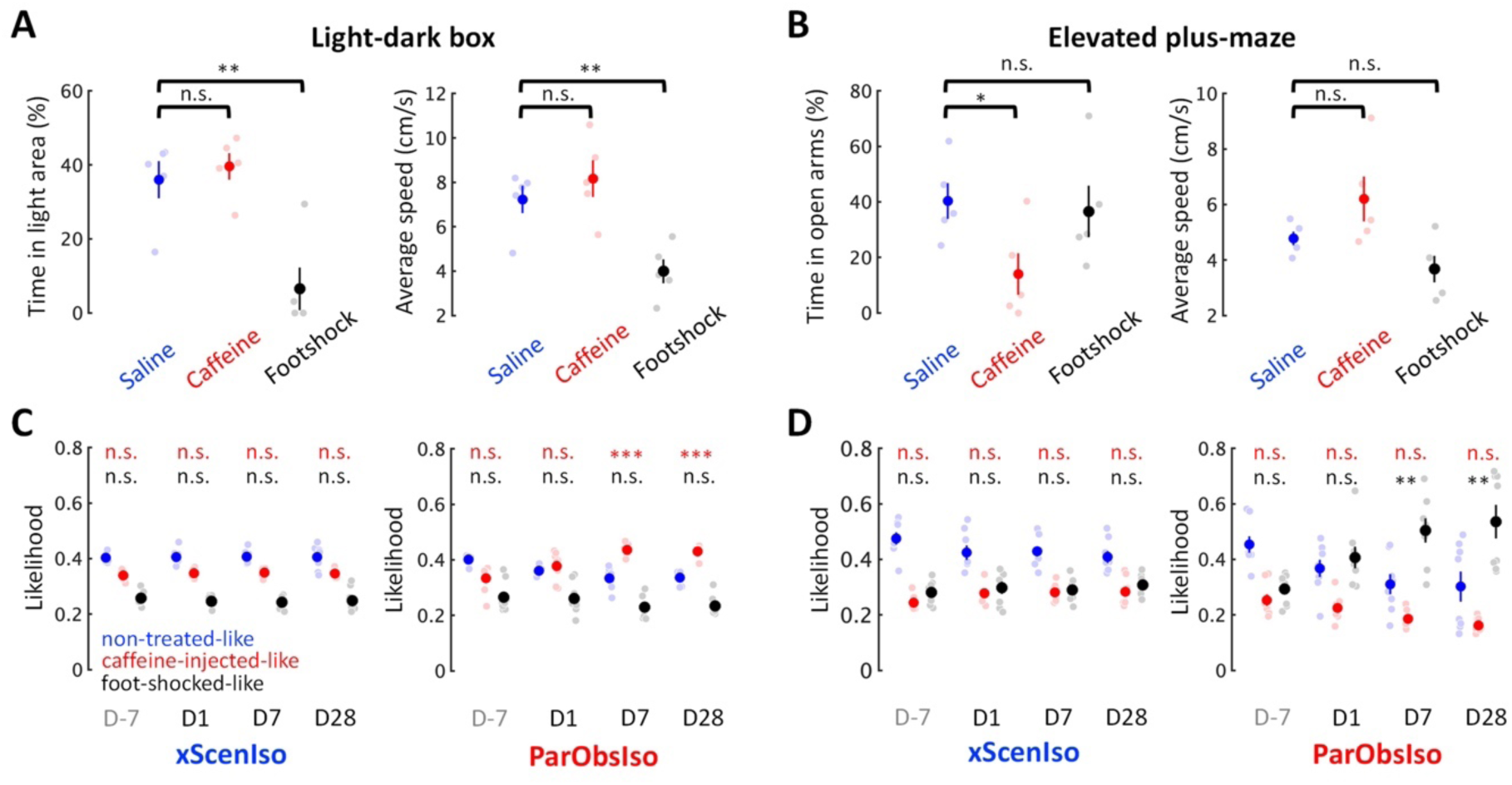
Comparison of behavioral characteristics in high-dimensional state-space indicates chronic somatic and cognitive anxiety developed in stress incubation. **(A)** Foot-shocked mice displayed less time spent in the light area and slower locomotion compared with the saline-injected and caffeine-injected mice in the light-dark box test. **(B)** Caffeine-injected mice displayed less time spent in the opened arms compared with the saline-injected and foot-shocked mice in the elevated plus-maze test. **(C)** In the light-dark box test, xScenIso mice stably showed non-treated-like behavioral characteristics after separated with its pair-housed partners, while ParObsIso mice increased their caffeine-injected-like behavioral characteristics in the corresponding period. **(D)** In the elevated plus-maze test, xScenIso mice kept showing highest likelihood of behavioral characteristics as non-treated-like after separated with its pair-housed partners, while ParObsIso mice increased their foot-shocked-like behavioral characteristics in the corresponding period. Note that comparison of behavioral characteristics in high-dimensional state-space was tested by one-tailed, two-sample Student’s t-test.

### Different developmental components were untangled from a single behavioral test

Through our fine-scale behavioral analyses, we further observed that, in the light-dark box and elevated plus-maze tests, key incubation features were consistently more obvious within stressor-free zones (dark area and closed arms) than within stressor zones (light area and open arms). To quantitatively investigate the differences between zones, we analyzed locomotor speed, as an independent behavioral index of mouse position, separately in stressor-free and stressor zones (Figure 6). As expected, two different behavioral patterns developed in the two zones: While speed differences consistently increased in stressor-free zones (Figure 6, B and D), speed differences in stressor zones only showed acute increases on Day1 (Figure 6, A and C). These results suggest that different psychological components may be measured in the two zones.

**Figure 6.**
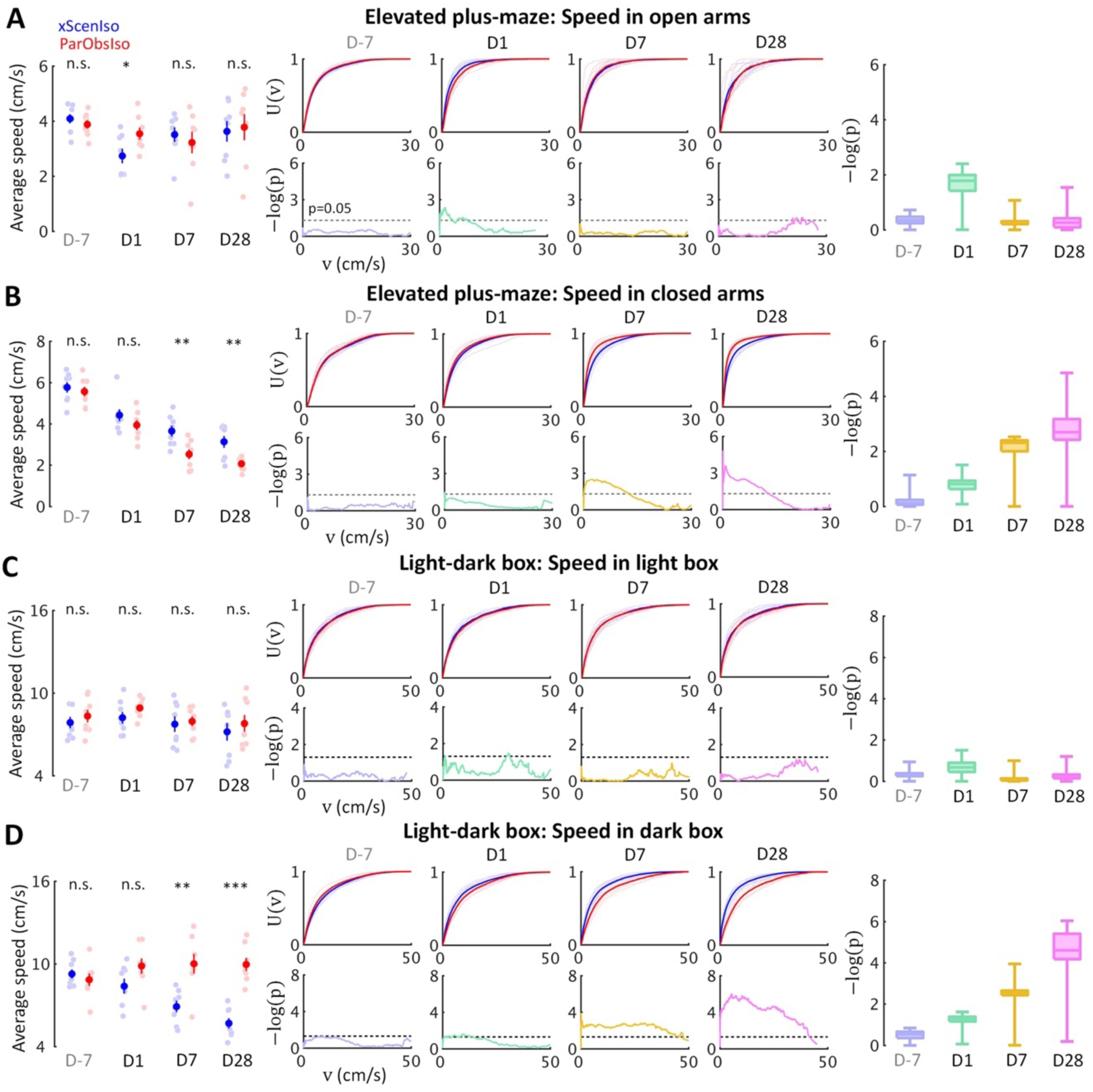
Gradually increasing anxiety and acute fear reaction were untangled in standard behavioral tests as different psychological components with distinctive developments. Locomotion speed shows acute difference only in stressor zones [open arms **(A)** and light area **(C)**], but incubated differences in stressor-free zones [dark area **(B)** and closed arms **(D)**].

Fear is a response to a known threat with a magnitude that increases with the strength of the threat, whereas anxiety is a response to uncertainty with a magnitude that increases with the uncertainty of a situation (Grupe and Nitschke, 2013). To better understand if these two developmental patterns observed under and close to the stressor may be uncertainty-related, we examined mouse locomotor speed in relatively uncertain and secure environments separately, using the open field test (Figure 7, A to C; n = 8 mice for each group) and the locomotor activity test (Figure 7D; n = 8 mice for each group). In the open field test, mice were exposed to a dimly lit environment with a gradient of spatial uncertainty from the field center (high uncertainty) to the field boundaries (low uncertainty). In the locomotor activity test, mice were habituated in a dark and smaller environment with familiar cues of bedding materials from their homecages indicative of less environmental stress. Compared to xScenIso mice, ParObsIso mice moved faster in the center region of the open field (Figure 7A), which was not observed in the periphery (Figure 7B). The difference between groups increased gradually towards the center but not in the periphery (Figure 7, A and B). The avoidance of spatial uncertainty by ParObsIso mice was also reflected in the shorter time they spent near the center (Figure 7C, left panel) and showed a shorter latency to the first rearing during the test (Figure 7C, right panel). In contrast, ParObsIso mice moved significantly slower on Day1 in the locomotor activity test and recovered on Days 7 and 28 (Figure 7D), suggesting an acute post-traumatic impact under low stress condition. These results suggest a possibility that the differences of uncertainty-related spontaneous behaviors incubate, whereas that of uncertainty-unrelated spontaneous behaviors attenuate.

**Figure 7.**
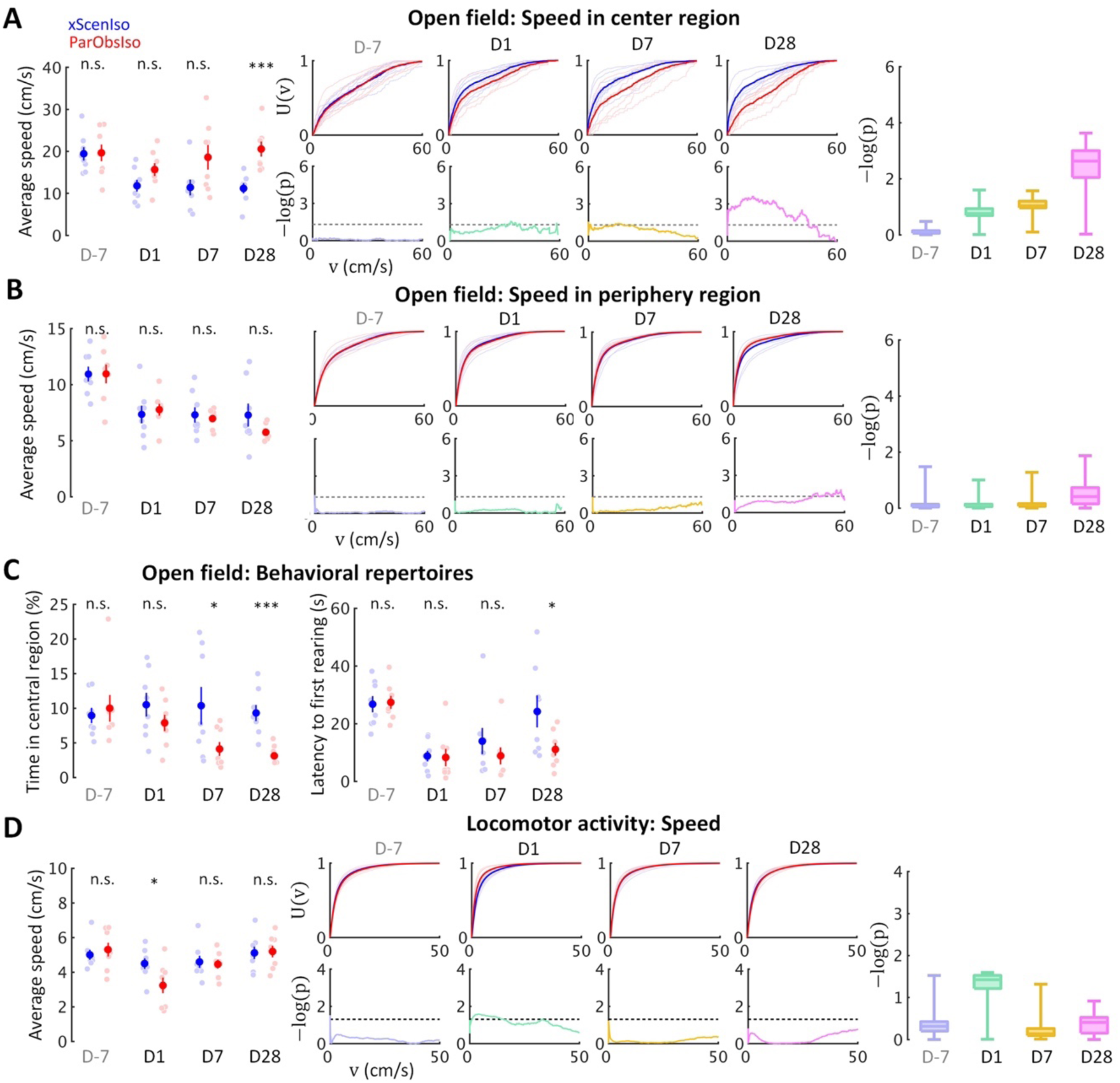
Behavioral testing in the open field test and locomotor activity test confirms distinctive psychological substrates and their corresponding development patterns. **(A)** Anxiety is evident in the open field test through a delayed onset of locomotor speed differences in the center region with higher spatial uncertainty. **(B)** The differences of locomotor speed observed in the center region did not occur in the periphery region with lower spatial uncertainty. **(C)** Less time spent in the central region by ParObsIso mice suggests their avoidance of a region with high special uncertainty, while shorter latency to their first rearing indicates their higher exploratory motivation. **(D)** In the locomotor activity test without stressors, acute effects of activity reduction recovered in the later post-traumatic period.

### Social differences weakly depend on emotional differences and vice versa

Emotional responses simplify and speed up animal reactions to complex external cues and are critical in corresponding social interactions (Anderson, 2016). To test potential changes of social interactions in ParObsIso mice, we examined mouse social motivation in a two-session social test (Figure 8A) where a non-social session was followed by a social session (n = 5 mice for each group). The social stimulus was either a female or a male stranger mouse. During the social session, ParObsIso mice spent less time in social approaches of nose poking toward both female and male strangers, starting from Day1, and remained less social compared to xScenIso mice during the post-traumatic period (Figure 8B). The time they spent in the interaction zone around the social target, however, did not differ significantly from that of xScenIso mice (Figure 8C; Supplemental Video 7). Both, ParObsIso and xScenIso mice, spent only a short but similar time on nose poking during the non-social session through the recordings on different days, with no significant difference between ParObsIso and xScenIso populations (Figure 8D), indicating that the observed difference of nose poking time was specific to social behavior. In addition, less social vocalization was recorded during the female stranger test for ParObsIso mice (Figure 8E). These observations showed that trauma induction also caused long-term differences in social interactions.

**Figure 8.**
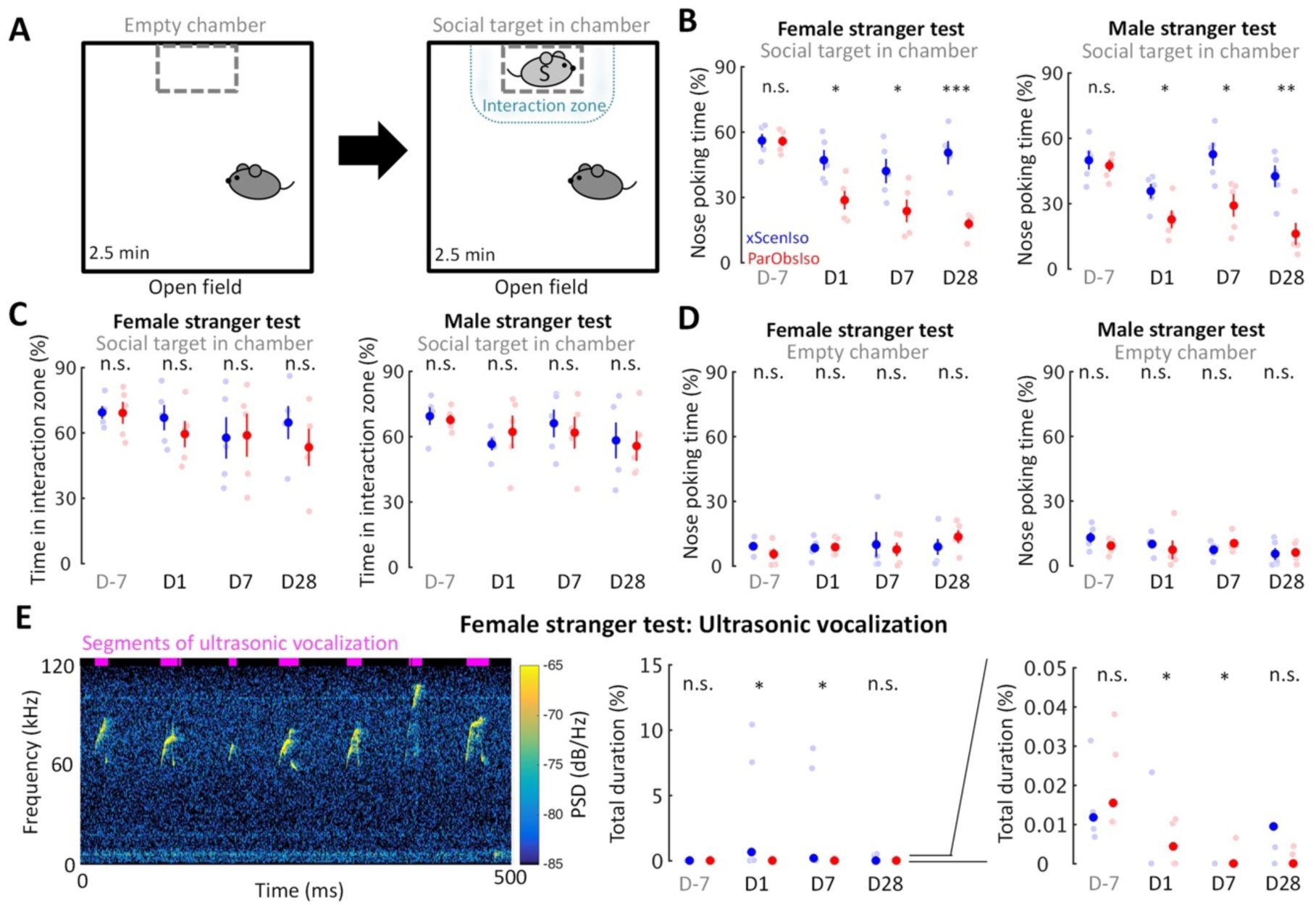
Acute psychosocial trauma decreases social interest. **(A)** The active social interaction test with consecutive non-social and social phases. During the social phase, motivation for social contact toward a stranger mouse (light gray S) was evaluated as the time spent for social approaches of nose poking and the time spent in the delineated interaction zone. **(B)** ParObsIso mice made fewer nose poking to both female (left) and male (right) strangers. **(C)** There was no significant difference in the time spent in the interaction zone during the social phase, suggesting a decrease of social interest instead of an active social avoidance. **(D)** There was no significant difference in the time spent of nose poking during the non-social phase, confirming that the observed differences of nose poking time stemmed from a specifically social root. **(E)** Spectrogram of short but conspicuous ultrasonic vocalizations emphasizes a specific behavioral repertoire during the social session in the female stranger test of a xScenIso mouse on Day1. More vocalization was recorded from xScenIso mice than ParObsIso mice on Days 1 and 7. Reduced ultrasonic vocalization during the social session of the female stranger test in ParObsIso mice attests to diminished social communication. Note that data points greater than 0.05% are not visible in the right panel which zoom in the data of the middle panel to emphasize the data distributions in the range of 0–0.05%. Data points and median, one-tailed Mann-Whitney U test; PSD, power spectral density.

To address the relation between emotion and social cognition, we next examined how the change of a particular social experience after the trauma induction could alter behavioral development of both individual spontaneous behaviors and social interactions. An important condition in our behavioral paradigm was the forced social isolation after trauma, of which the potential effects should further be identified. We examined the impact of post-traumatic social condition on behavioral developments by the second control group of mice [Partner-Observing-Partner-Pair-Housed (ParObsParPH) mice; Figure 1C] as each of them was kept pair-housed with its attacked partner after trauma induction. Noteworthily, developments of the differences in spontaneous behaviors did not occur in ParObsParPH mice (Figure 9A, upper panels; Supplemental Figure 5A), but importantly, developments of behavioral differences in social interactions showed the differences of ParObsIso mice (Figure 9B, left panel). This observation implies the sufficiency of socially isolated condition for differences of individual spontaneous behaviors but not for that of social interactions, suggesting a weak inter-relationship between the development of emotional differences and that of social differences.

**Figure 9.**
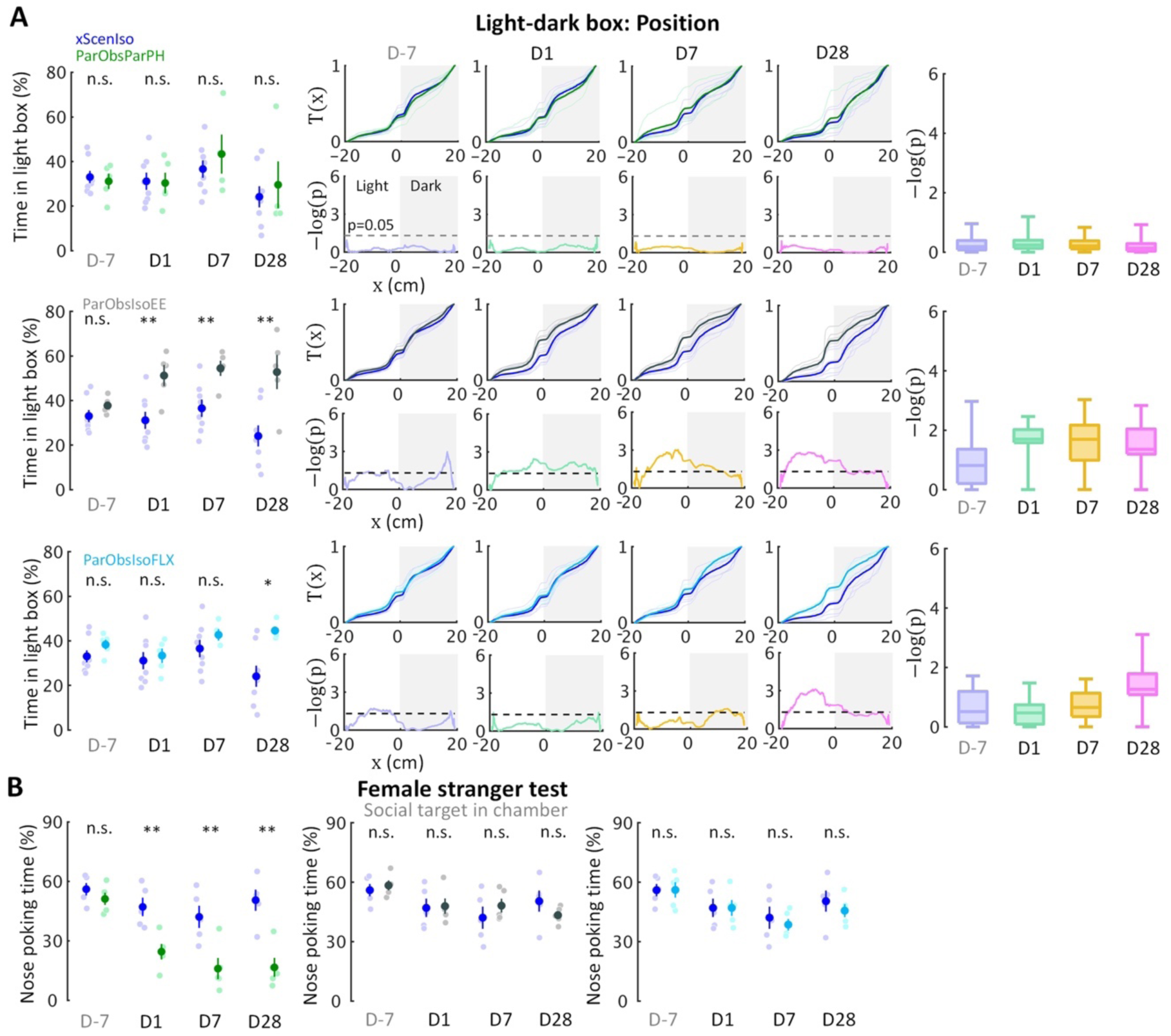
Developments of emotional and social differences are not inter-dependent. **(A)** Positions in the light-dark box test indicate that ParObsParPH mice did not develop the chronic stress reactions, ParObsIsoEE mice developed chronic stress reactions which were stronger than that of ParObsIso mice in the early phase, and ParObsIsoFLX developed stress reactions in the late phase. **(B)** Nose poking times in the female stranger test indicate that ParObsParPH, but not ParObsIsoEE and ParObsIsoFLX, mice develop the social differences of ParObsIso mice.

ParObsParPH mice experienced the same scenario of trauma induction as ParObsIso mice but had different post-traumatic social experiences, leading to a partial rescue of differences in spontaneous behaviors. To test the potential significance of the post-traumatic social factor in behavioral developments, we conducted two additional control experiments by alternating the post-traumatic environmental and physiological conditions, respectively, with the third control group of mice where each of them was provided with toys during social isolation after trauma induction [Partner-Observing-Isolated-Environment-Enriched (ParObsIsoEE) mice; Figure 1D], and the fourth control group of mice [Partner-Observing-Isolated-Fluoxetine-Treated (ParObsIsoFLX) mice; Figure 1E] as each of them was injected with fluoxetine daily after trauma. ParObsIsoEE mice showed similar nose poking times in the female stranger test as xScenIso mice (Figure 9B, middle panel); however, they showed the behavioral differences of ParObsIso mice in the light-dark box test, which even had a stronger difference starting from the early phase (Figure 9A, middle panels). For ParObsIsoFLX mice, while they showed less behavioral difference during the early post-traumatic phase in the light-dark box test (Figure 9A, bottom panels) and did not display a reduction of nose poking times in the female stranger test (Figure 9B, right panel), their behavioral difference in the light-dark box test reached significance in the late phase and the behavioral difference in elevated plus-maze test was more obvious in the closed arms after trauma induction (Figure 9A, bottom panels; Supplemental Figure 6A). The partial recusing and even partial strengthening of behavioral differences observed from these experimental alterations of post-traumatic social and non-social factors emphasize the sensitivity and complexity of context-wide behavioral development over stress incubation.

### Social relationship determined context-wide behavioral development over stress incubation

To address the psychobehavioral basis of stress incubation underlying the complexity of context-wide behavioral developments that may be weakly inter-dependent, we conducted the fifth control experiment where each of the mice was kept pair-housed with a stranger after trauma induction [Partner-Observing-Stranger-Pair-Housed (ParObsStrPH) mice; Figure 1F], altering the post-traumatic social relationship in the ParObsParPH paradigm. In the ParObsStrPH paradigm, if the pair-housed strangers were the socially defeated intruders of the trauma inductions, we observed aggressive attacks toward strangers by all focal mice (n=5 out of 5) during their pair-housing after trauma. Similar aggression was observed among ParObsIso mice, but not toward their defeated partners, when they were group-housed after trauma induction. Because of these observations of partnership-dependent aggressive or non-aggressive behaviors, we only conducted in-depth behavioral tests with the ParObsStrPH mice pair-housed with non-defeated strangers and recorded their behavior. ParObsStrPH mice did not display the development of behavioral differences in the light-dark box, elevated plus-maze, and female stranger tests (Figure 10A, upper panels; 10B, left panel; Supplemental Figure 6B). Only their position in the light-dark box test showed an acute difference (Figure 10A, upper panels). These results indicate that a single factor of social relationship after trauma induction may govern context-wide developments into long-term behavioral differences.

Following the evidence that social relationship governs the behavioral development after the trauma induction, we further examined potential contribution of social experience on trauma induction by conducting two additional control experiments with the alternation of social factors during trauma induction. In the sixth control group of mice [Non-Aggressor-Exposed-Isolated (xAggrExpIso) mice; Figure 1G], the aggressor mice for trauma induction were replaced by non-aggressive strangers. xAggrExpIso mice did not show the developments of behavioral differences in both spontaneous behaviors (Figure 10A, bottom panels) and social interactions (Figure 10B, right panel), confirming that, instead of the increased number of mice during trauma induction (visual exposure to 5 different aggressors), social aggression is necessary to induce development of behavioral differences.

For the seventh group of mice [Stranger-Observing-Isolated (StrObsIso) mice; Figure 1H], each of the mice observed different stranger mice being attacked by different aggressors and stayed together with each stranger between aggressive encounters before isolation. StrObsIso mice did not experience any physical stress from strangers or aggressors. During trauma induction, two notable behavioral differences were observed: While tail rattling during aggressive encounters and hiding under bedding material with the partner during resting were observed in 83% (n = 39 out of 47 mice) and 100% (n = 47 out of 47 mice) ParObsIso mice (Supplemental Video 8), respectively, no such behaviors were shown by StrObsIso mice (n = 0 out of 20 mice; n = 0 out of 20 mice). In ParObsIso mice, the frequency of tail rattles dropped progressively during aggressive encounters (Figure 11A), representing a transient reaction during trauma induction. These results suggest that social relationship may rapidly modulate emotional impact and social reactions during the trauma induction. Moreover, chronic behavioral differences were not observed in StrObsIso mice (Figure 11, B to F), which further excluded potential effects from salient, non-specific environmental manipulation (e.g. rotation through aggressors’ home cages) and sensory shock (e.g. olfactory cues from urine and vocalization indicating fear) during trauma induction. Taken together, these results show that social relationship constitutes as a critical factor of trauma induction and its following context-wide developments of behavioral differences.

Considering social relationship as the key factor, an important question remained: Why did behavioral differences in the putatively uncertainty-related component of spontaneous behaviors gradually increase during stress incubation, but not remain constant or attenuate? To address this issue, we examined the persistence of social memory in a partner-revisiting test on Day28 (Figure 12A; n = 5 mice for each group). Social stimuli were the previous partner and a stranger mouse, both immobilized to allow enough social cues to be attractive, but no active interaction with the focal mouse. To avoid possible influences of social cues from socially defeated mice, stranger mice used to test xScenIso and StrObsIso mice were partners of ParObsIso mice. Strikingly, ParObsIso mice as well as ParObsIsoPH mice, which were separated from their partners right before their partners got immobilized for the tests, spent three times as much time allogrooming or pushing their previous partners as did xScenIso, xAggrExpIso, and StrObsIso mice (Figure 12B and Supplemental Video 9). The preference of ParObsIso mice to their previous partners, together with the contribution of social relationship to context-wide developments of behavioral differences after separation, implies that the uncertainty of partnership may be a key mechanism of the gradually increasing difference in uncertainty-related behaviors in our paradigm.

## Discussion

We present an approach to interpret otherwise challenging and inconclusive behavioral data and use it to study stress incubation in laboratory mice. The results demonstrate a system-level view of experimentally disentangled components, processes, and determinants in stress development. We report the asymmetry of brain-wide microstructural changes and the strengthening of an ACC-centered network in mice after acute witnessing social stress. Based on the context-wide observations from our experiments, we propose that social relationship, as the single common factor, may underlie the otherwise independent development of stress incubation. Our study provides technical and conceptual advances which could be considered in the study of human psychiatry disorders such as PTSD.

### Detection and identification of animal emotion

We argue that classical behavioral analyses of standard tests can be too coarse to capture intricate emotional states. The traditional approach to identify animal emotion is to test if animals show particular behaviors specified by the experimental test assumed when it was designed (Walf and Frye, 2007). However, experimental animals normally display obvious but not inter-supporting behaviors in different tests with the same logic and assumptions (Ramos, 2008). Furthermore, even if the behavioral results are consistent with the expectation, they can still be alternatively explained (Garcia et al., 2008). This ambiguity reaches deeply into the history of widely used behavioral tests and therefore have resulted in a considerable amount of inconclusive and seemingly paradoxical results, which are usually left for discussion or remain unreported (Carobrez and Bertoglio, 2005; Crusio, 2013; Engin and Treit, 2008; Ennaceur, 2014; Hascoët et al., 2001; Henriques-Alves and Queiroz, 2016; Kulesskaya and Voikar, 2014). The limitations can be due to circular arguments embedded in a reductive logic. Classically, researchers addressed the difficulties by an effort to show a proof-of-concept (Vasconcelos et al., 2012). Therefore, studies frequently use multiple tests to assess the same psychological phenomenon. With this approach, research on animal emotion usually emphasize their ability to show human behavioral and physiological conditions (face validity), to reproduce pharmacological effects in human (predictive validity), and to share the same biological processes as human (construct validity) (Calhoon and Tye, 2015), although a few studies have taken the possibility of human-unique, animal-unique, and human-animal-sharing emotions under consideration (Anderson and Adolphs, 2014). To date, for the long and widely used behavioral tests, a standardized method objectively identifies and quantifies emotion from a continuous, high-dimensional state-space of behaviors is still absent.

In this study, we introduced fine-scale behavioral analysis and state-space behavioral characterization to access animal emotion from standard behavioral tests which give inconclusive results when analyzed and interpreted in the traditional way. Taking our observations as an example, higher nest wall (Figure 2A), higher baseline corticosterone levels (Figure 2C), more time spent in the far end of closed arms (Figure 4A), less exploration from closed arms to center (Figure 4C, left panel), longer freezing time (Figure 4C, right panel), and slower locomotion (Figure 4B) in the elevated plus-maze test, more time spent in the center region (Figure 7C, left panel), and shorter latency to the first rearing (Figure 7C, right panel) in the open field test, slower locomotion in the locomotor activity test (Figure 7D), and less nose poking (Figure 8B) and vocalization (Figure 8E) in the stranger tests are classically more acceptable as stress reactions. However, we also note that increased body mass (Figure 2B), more time spent in light area (Figure 3A), faster locomotion (Figure 3B), more transfers (Figure 3B, right panel), shorter latency to the first transfer (Figure 3C, right panel) in the light-dark box test, spending similar time around social targets in the stranger tests (Figure 8C), and more social reactions to previously pair-housed partners (Figure 12B) are more controversial in classic readouts. As an example, if only focusing on the observations of increased body mass and higher nest wall (Figure 2, A and B), they can be interpreted as having better emotional stability and therefore nicer nests and better appetites, or manifesting protective responses of anxiety by hiding behind higher nest walls and engaging in anxiety-induced binge eating (Goto et al., 2014; Otabi et al., 2017). Discussing potential interpretations or possible factors of these massive observations can be endless and easily lead to skeptical arguments especially if the focused psychobehavioral substrate is complex. With both richer measurements and quantitative analyses, we were able to discover subtle behavioral differences and identify otherwise obscured behavioral details in stress incubation. In addition, we propose to interpret psychological meaning based on experimental comparison and correlation with physical variables of the testing environment, rather than based on the expectation of presumed observations, as traditionally done. Importantly, founded on computational ethology (Anderson and Perona, 2014; Chen et al., 2013; Nath et al., 2019), providing a validly consistent overview of data interpretations for context-wide observations across the experiments becomes possibly more reliable and more effective.

### From stress incubation in mice to human PTSD development

Behavioral paradigms of laboratory rodents that simulate PTSD were established to expose the mechanistic insights of long-term fear memory following acute physical stress (Balogh et al., 2002; Philbert et al., 2011) or learned depression after repeated social defeat (Sial et al., 2016; Warren et al., 2013). The diversity of PTSD models in mice is even highlighted by a paradigm of trauma-free pharmacologically-induced memory impairments in mice, which was recognized as a PTSD model and further identified the corresponding pathophysiological mechanism (Kaouane et al., 2012). Although witnessing social defeat models in rodents were developed, an identification of post-traumatic stress incubation was still challenged by significant effects of the prolonged peri-traumatic stress development from the repeated trauma induction for more than a week (Patki et al., 2014; Warren et al., 2013).

In this study, we found little support for the common view that an association among psychological states, such as emotion and social motivation, govern the developmental process (Andrews et al., 2007; Bryant et al., 2017; DSM-5, 2013; DSM-III, 1980; Ehlers and Clark, 2000; Hayes et al., 2012; Pamplona et al., 2011; Schnyder and Cloitre, 2015; Siegmund and Wotjak, 2006; Zoladz and Diamond, 2013). We observed continuous growth of the differences in uncertainty-related spontaneous behaviors while that of uncertainty-unrelated spontaneous behaviors had long vanished (Figures 4 to 7). The time course of substantial social differences also led the substantial differences of uncertainty-related spontaneous behaviors (Figures 4, 5, and 8). In addition, pair-housing with partner mice selectively rescued the increased differences of spontaneous behaviors but not social differences (Figure 11), and, in contrast, environmental enhancement selectively strengthen the differences of spontaneous behaviors but reduced the differences in social behavior (Figure 9A, middle panels). However, even with this weak behavioral correlate of different psychological aspects in stress incubation, we found that there is a single factor, social relationship, commonly mediating diverse behavioral developments. Alternation of a single social cognitive factor, social relationship, eliminated behavioral differences from mice without traumatic experience (Figures 10 and 11). Conceptualization of social support as a “stress buffer” have been proposed to explain the positive association between responsive social resources in a small social network and adverse effects of stressful events (Cohen and Wills, 1985). Indeed, among all rescue controls, including social, environmental, or pharmacological approaches (Figure 9, C to F), pair-housing with non-defeated stranger mice showed the best rescuing effects on diverse behavioral developments in our paradigm (Figure 12). In the collective model (Figure 13A), behaviors slowly change and influence each other during stress incubation until a new, abnormal equilibrium is reached; this new equilibrium is then defined as PTSD. However, based on our observations and tests, we propose that a specific internal cause and its related processes play a dominant role in stress incubation. In this unitary model (Figure 13B), a single common factor underlies the otherwise independent development of post-traumatic behaviors in mice.

**Figure 10.**
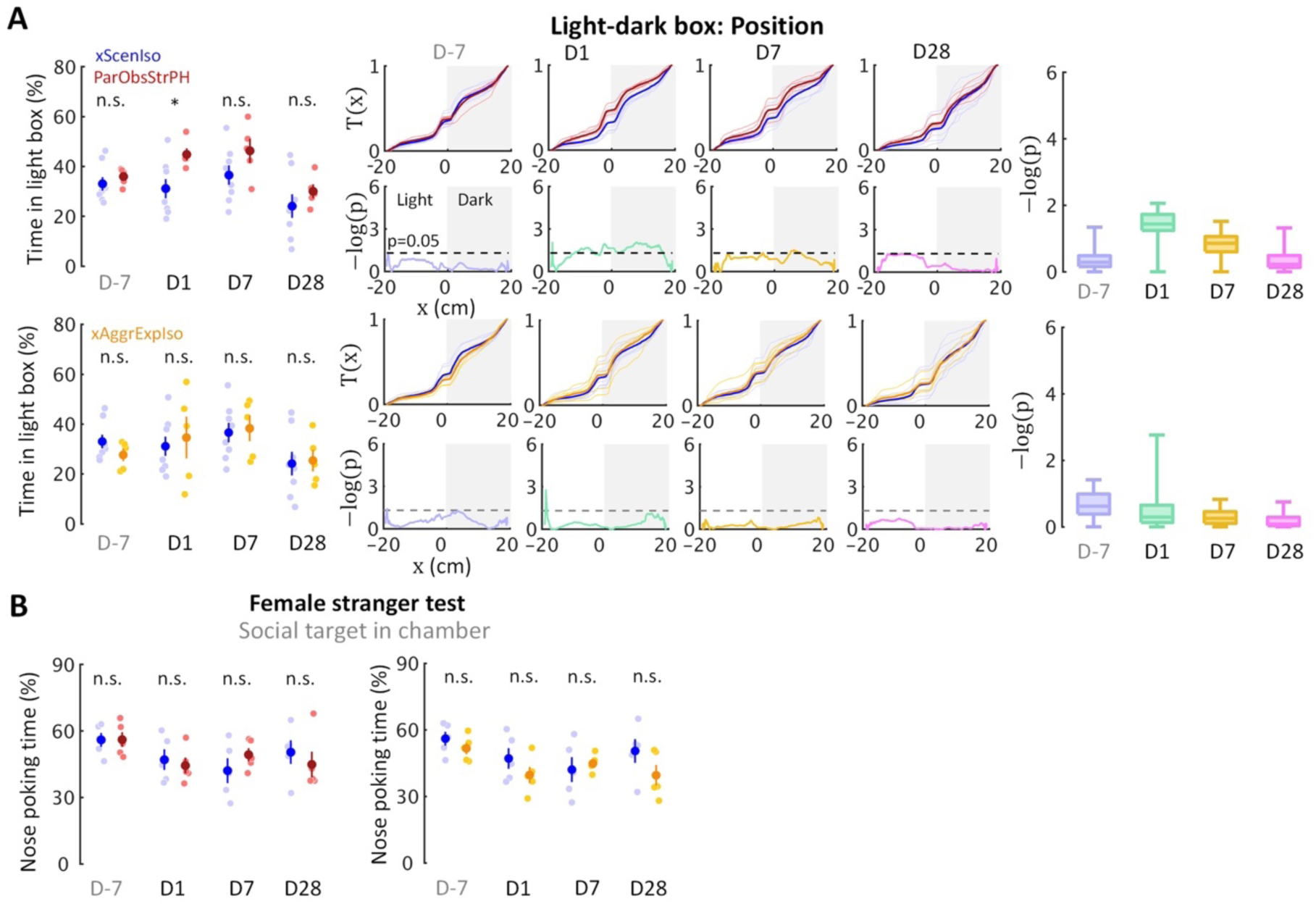
Control experiments identify necessity of relationship-dependent vicarious defeat for anxiety incubation. **(A)** Positions in the light-dark box test indicate that both xAggrExpIso and ParObsStrPH mice did not develop the behavioral differences in the late phase, although ParObsStrPH mice displayed the difference in the early phase. **(B)** Nose poking times in the female stranger test indicate that xAggrExpIso and ParObsStrPH mice did not develop the social differences.

**Figure 11.**
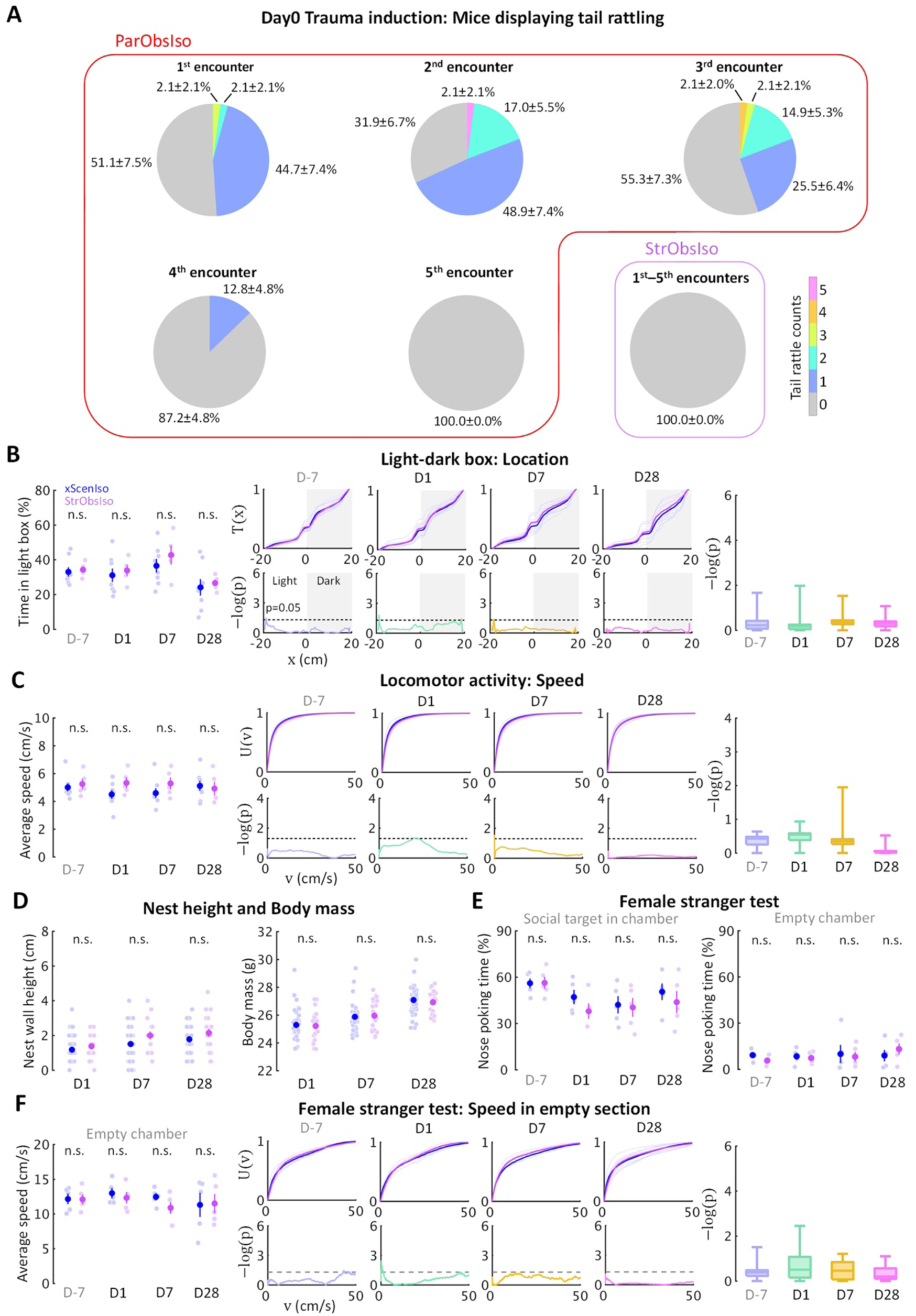
Social relationship in trauma induction determines stress development. **(A)** Social relationship increased emotional impact, evidenced by tail rattling behavior during trauma induction. **(B–F)** No significant acute or chronic difference was found in StrObsIso mice.

**Figure 12.**
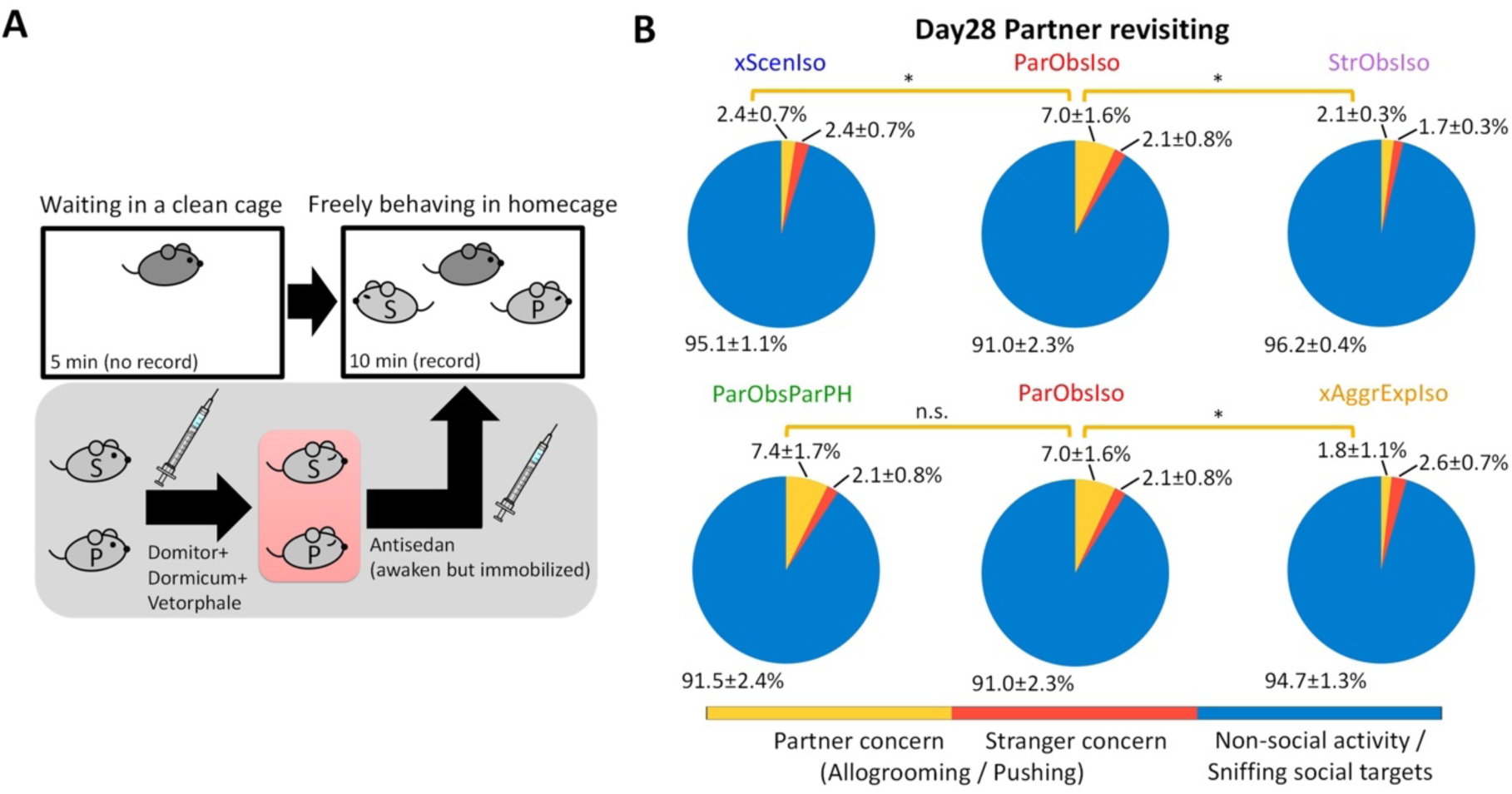
Long-term memory of partnership correlates with anxiety incubation. **(A)** The partner-revisiting test. A stranger mouse (light gray S) and the previously pair-housed partner (light gray P), both immobilized, were presented as social targets. Pink rectangle, heating pad. **(B)** ParObsIso and ParObsParPH mice showed significantly longer allogrooming or pushing their partners (yellow, % of time spent in partner concern behavior) than either xScenIso, xAggrExpIso, or StrObsIso mice. Standard errors were calculated from bootstrapped data.

**Figure 13.**
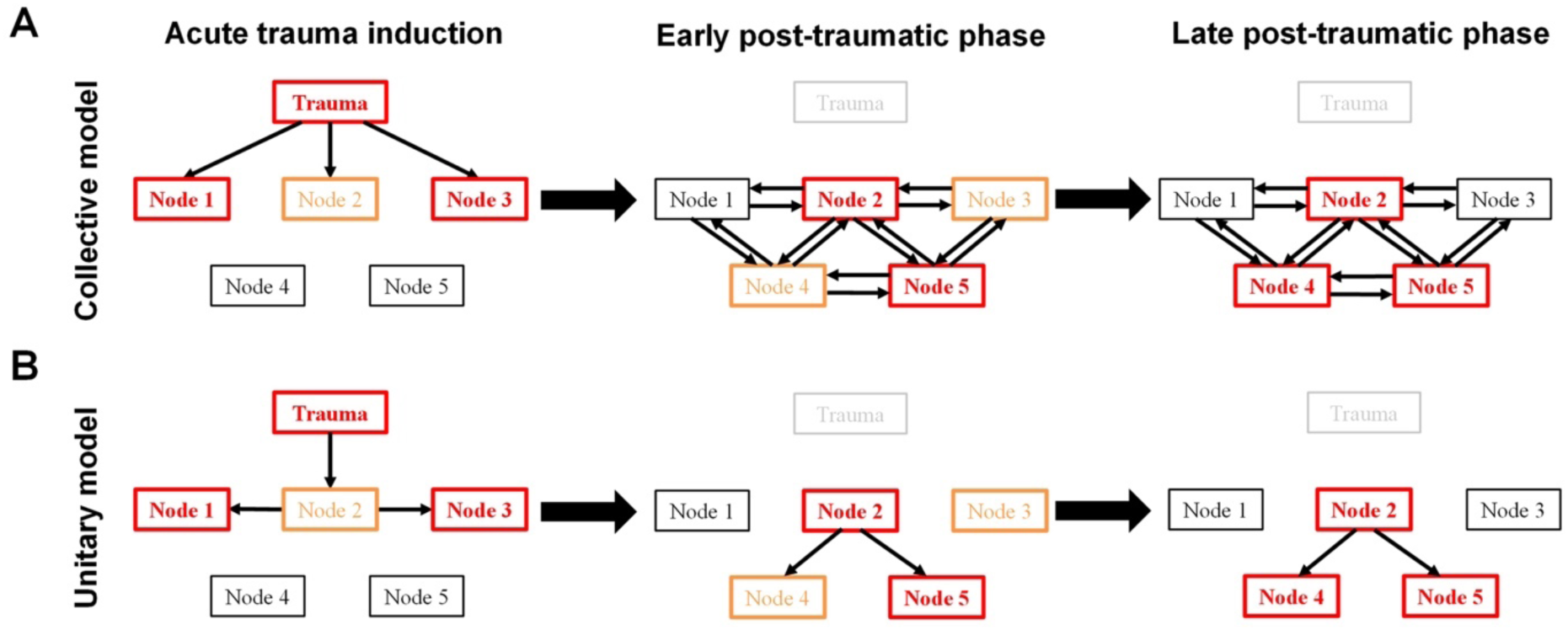
Two conceptual models of the post-traumatic stress incubation process. **(A)** In the collective model, different psychological elements (Node #, a type of emotion or cognition) influence each other during different phases of stress incubation. **(B)** In the unitary model a single common factor underlies the development of post-traumatic behaviors. Orange and red boxes represent medium and strong trauma-induced differences, respectively. It is assumed that each psychological element can be connected to an associated neural substrate with trauma-induced dynamic changes.

From DTI scanning, we also found asymmetric microstructure differences and an asymmetric enhanced, ACC-centered network in the two brain hemispheres, 28 days after the witnessing of social stress. This is in agreement with previous finding that the right but not the left ACC controls observational fear learning in mice (Kim et al., 2012). The brain heavily integrates not only external but also internal causes (Donaldson et al., 2015; Funamizu et al., 2016; Kohl et al., 2018; Larkum, 2013; Lee et al., 2014; R. X. Lee et al., 2015; Matias et al., 2017; Murugan et al., 2017; Remedios et al., 2017; Roome and Kuhn, 2018; Tononi et al., 2016; Zelikowsky et al., 2018). The potentially slow and global change of brain dynamics may arise from an altered dynamic in social bonding circuits through their interconnected nodes (Ko, 2017). By correlating with freezing behavior following a single scrambled foot shock in mice, early inhibition of PTH2R (parathyroid hormone 2 receptor)-mediated TIP39 (tuberoinfundibular peptide of 39 residues) signaling in the medial amygdalar nucleus was demonstrated to enhance fear memory much later (Tsuda et al., 2015). Following this line, a delay in generalized avoidance was proposed developing from an amplification of fear expression (Houston et al., 1999; Pamplona et al., 2011; Sillivan et al., 2017). Interestingly, the connections between the piriform and perirhinal cortices decreased (Figure 2, H and I). The piriform and perirhinal cortices are the two core parahippocampal structures involve in the kindling phenomenon, the daily progressive increase in response severity of both electrographic and behavioral seizure activity (McIntyre, 2006; McIntyre and Kelly, 2006) supposedly linked to fear conditioning in rat PTSD models (Knox et al., 2012; Rau et al., 2005). In human PTSD research, the amygdala, medial prefrontal cortex, and hippocampus are the brain regions traditionally focused on (Shin et al., 2006), with reports emphasizing the morphology of the right hippocampus (Gilbertson et al., 2002; Pavić et al., 2007). Our findings extend these observations to laboratory rodent model of witnessing stress under experimental conditions.

The medial prefrontal cortex (mPFC), which includes ACC and prelimbic cortex (PL) in mice, have been reported to exhibit both functional and physiological asymmetry between hemispheres. For examples, the right mPFC was reported to control the acquisition of stress during hazardous experiences while the left mPFC was found to play a dominant role in translating stress into social behavior (E. Lee et al., 2015). The effects of erythropoietin on inhibitory synaptic transmission in the left and right PL of mice were also found to be opposite (Dik et al., 2018). Furthermore, neuromodulatory systems can play an important role in lateralized circuitry processing. Oxytocin receptor expression is lateralized as there are more OXTR-2 receptors on the left side of the auditory cortex in adult females (Marlin and Froemke, 2017). Based on this asymmetric nature, an oxytocin-mediated balancing of left and right cortical synaptic inhibition was reported to enable maternal behavior in mice (Marlin et al., 2015). Stress-induced mesocortical dopamine activation was found for the right mPFC but not the left (Sullivan and Gratton, 1998). Additionally, serotonin selectively regulates mPFC callosal projection neurons (Avesar and Gulledge, 2012; Stephens et al., 2014), suggesting specific roles of the communication between left and right mPFC although they are functionally distinct. These lateralized changes on the right side due to stress experience is consistent with our observation of the changes of cortical microstructures and fibers on the right hemisphere in mice showing PTSD-like behavior. This is comparable to the predominantly right cortical volumetric differences in human PTSD (Bremner et al., 2005; Gilbertson et al., 2002). While most mechanistic or neuronal recording studies in mice only focused on one side of the brain or simultaneously regulated both sides of the brain, our data highlight a requirement for special attention on hemisphere-specific control of cognitive and emotional processes.

Although we focused on stress incubation in mice, our work may provide several important insights to complex human psychiatry, especially PTSD development. Complex developmental trajectories of human PTSD symptoms were demonstrated with associated physiological and environmental regulators (Bryant et al., 2013), yet the combination of psychological therapy and pharmacotherapy does not provide a more efficacious treatment than psychological therapy alone (Hetrick et al., 2010; Mataix-Cols et al., 2017). Our research provides a foundation to test pharmacotherapies on a system-level that integrates multiple mechanisms underlying highly diverse behavioral consequences. The efficacy of pharmacotherapies could therefore be improved. While much remains to be considered before clinical applications, our study establishes a solid basis to uncover psychological theories for therapeutic strategies (Schnyder and Cloitre, 2015). Compared with the classic view of PTSD development as a process of complex associations (McFarlane, 2010), we propose to consider human PTSD development as a process with unitary origin. While the core factor in a unitary model of human PTSD is not necessarily social bonding, we suggest a special focus on affective bonding as the core factor for patient diagnosed with witnessing PTSD. We also expect that behavioral signs of PTSD development could be detected in humans already shortly after the traumatic event. The fine-scale behavioral analyses we introduced here provides a simple, non-invasive analytic tool to capture informative behavioral details and is not limited to laboratory animals. It opens a new window for early detection and prediction, with the potential to prevent the development into PTSD.

### Limitations of the Study

Although we have conducted an in-depth investigation covering multiple dimensions of behavioral phenotype, our tests and paradigms only focus on a small part of all the possibilities of an animal facing the great uncertainty in nature and achieving countless tasks through its live. In addition, since our data did not replicate common findings of social avoidance followed by social defeat stress, a “dose response” of observational stress could govern this “micro-defeat” which did not result in a “standard constellation” of PTSD-like behaviors in mice. Finally, individual performance across different tests and the corresponding cross-testing measurement effects on stress development, as a factor may be prone to systematic errors, was ruled out from our study design and not examined.

## Materials and Methods

All animal experiments were approved by the Institutional Animal Care and Use Committee (IACUC) in the animal facility at the Okinawa Institute of Science and Technology (OIST) Graduate University, accredited by the Association for Assessment and Accreditation of Laboratory Animal Care (AAALAC). All animal procedures were conducted in accordance with guidelines of the OIST IACUC in the AAALAC-accredited facility.

### Study design

The goal of this work is to identify diverse psychological aspects, temporal patterns, and associations of behavioral development in mice after a single trauma induction. Pre-specified hypotheses stated that (i) development of behavioral differences is already represented in behavioral details during the early post-traumatic phase, while (ii) the behavioral differences of spontaneous behaviors and social interactions are inter-dependent. Our data support the first pre-specified hypothesis, but not the second. All other hypotheses were suggested after initiation of the data analyses.

We approached the research goal by developing a novel mouse model of psychosocial trauma under highly controlled conditions (psychosocial manipulations, subjective experiences, and genetic background) and by applying fine-scale analysis to standard behavioral tests. To induce acute witnessing trauma, a pair-housed mouse observed how its partner got bullied by a larger, aggressive mouse on the day of trauma induction. After this trauma, the observer mouse was isolated and developed behavioral differences compared to control mice in the ensuing weeks. The control groups included (i) mice isolated without experiencing trauma induction, (ii) mice isolated after observing how a stranger mouse got bullied by a larger, aggressive mouse, (iii) mice isolated after exposed to a non-aggressive stranger mouse, (iv) mice which were pair-housed with their defeat partners after observing how its partner got bullied by a larger, aggressive mouse, (v) mice which were pair-housed with strangers after observing how its partner got bullied by a larger, aggressive mouse, (vi) mice isolated with their environment enriched, and (vii) mice isolated with daily injections of fluoxetine. The behavioral tests included the light-dark box test, elevated plus-maze test, open field test, locomotor activity test, active social contact test to a female stranger, active social contact test to a male stranger, and partner-revisiting test. The non-behavioral tests included the body mass measurement, nest wall height measurement, baseline corticosterone concentration test, and *ex vivo* diffusion tensor imaging.

No statistical methods were used to predetermine sample sizes. Animal numbers were determined based on previous studies (Takahashi et al., 2015). 8-week-old male C57BL/6J mice, 16-week-old female C57BL/6J mice, and 20-week-old or older male Slc:ICR mice were used. No data were excluded. No outliers were defined. Mice were from different litters. Mice were randomly paired. A focal mouse was randomly selected from each pair of mice. Mice were randomly allocated into experimental groups. Testing order among groups was counterbalanced. Strangers and aggressors were randomly assigned. All behavioral tests were conducted in quintuplicate to octuplicate sampling replicates. All behavioral tests were conducted in single to quadrupole experimental cohorts. All other records were conducted in quadruplicate to septuplicate experimental cohorts. The investigator was blinded to behavioral outcomes but not to group allocation during data collection and/or analysis.

The endpoints were prospectively selected. Partner mice were expected to get minor injuries from aggressor mice during aggressive encounters; typically, attack bites on the dorsal side of posterior trunk (Takahashi et al., 2015). The aggressive encounter and all further experiments were terminated once (i) the partner mouse showed severe bleeding or ataxia, or (ii) the aggressor mice showed abnormal attack bites on any other body part. Partner mice fulfilling criteria (i) were euthanized. Aggressor mice fulfilling criteria (ii) were not used in any further experiments. If any aggressive sign (sideways threat, tail rattle, pursuit, and attack bite) was shown by the partner mouse, all further experiments with the partner mouse, aggressor mouse, and observer mouse were terminated.

### Overview

In total, 527 male C57BL/6J mice (CLEA Japan, Inc.), 49 female C57BL/6J mice (CLEA Japan, Inc.), and 33 male Slc:ICR mice (Japan SLC, Inc.; retired from used for breeding) were used in this study. In CLEA Japan, nursing females were individually housed (CL-0103-2; 165×234×118 mm), while pups were separated on P21 according to gender and housed ≤15 mice per cage (CL-0104-2; 206×317×125 mm). Pups were re-arranged on P28 according to their weights and housed ≤13 mice per cage (CL-0104-2). Mice were shipped in boxes each with 10 – 30 mice to the OIST Animal Facility. In the OIST Animal Facility, mice were housed in 380×180×160-mm transparent holding cages (Sealsafe Plus Mouse DGM - Digital Ready IVC; Tecniplast Inc., Quebec, Canada) bedded with 100% pulp (FUJ9298101; Oriental Yeast Co., Ltd., Tokyo, Japan) under a 12-hr dark/light cycle (350-lux white light) at a controlled temperature of 22.7 – 22.9 °C, humidity of 51 – 53%, and differential pressure of -14 – -8 Pa with food and water available ad libitum. Circadian time (CT) is defined to start at mid-light period and described in a 24-hr format, i.e. light off at CT 6:00.

The experimenter and caretakers wore laboratory jumpsuits, lab boots, latex gloves, face masks, and hair nets when handling mice and performing experiments. Handling of mice during the dark cycle was done under dim red light and mice were transported in a lightproof carrier within the animal facility. For mice in experimental and control groups tested on the same dates, the testing order was alternated. Surfaces of experimental apparatuses were wiped with 70% ethanol in water and dry paper tissues after testing each mouse to remove olfactory cues. Each mouse was only used for one behavioral test (in total 4 records with intervals of 6 – 21 days) to avoid confounded results due to cross-testing and to minimize measurement effects on its psychological development (Krishnan et al., 2007).

### Pre-traumatic period (Day-21 to Day0)

To establish partnerships between mice, a male C57BL/6J mouse (focal mouse; 8 weeks) was pair-housed with another male C57BL/6J mouse (partner mouse; 8 weeks) for 3 weeks (Day-21 to Day0, with trauma induction on Day0). The partner was initially marked by ear punching. The holding cage was replaced once per week, with the last change 3 days before the traumatic event (Day-3).

To establish the territory of an aggressor mouse in its homecage, an Slc:ICR mouse (aggressor mouse; ≥20 weeks) was pair-housed with a female C57BL/6J mouse (female mouse; 16 weeks) for 3 weeks (Day-21 to Day0). The holding cage was replaced with a clean one once a week, with the last change one week before the traumatic event (Day-7).

Aggression level of aggressors was screened on Days -5, -3, -1 through intruder encounters (Miczek and O’Donnell, 1978) toward different screening mice to determine appropriate aggressors to be used for trauma induction on Day0. Aggression screening was carried out in the behavior testing room at 22.4 – 23.0 °C, 53 – 58% humidity, -4 – -3 Pa differential pressure, and 57.1 dB(C) ambient noise level during the light period (CT 4:00 – 6:00) with 350-lux white light. After the female and pups with the aggressor were taken out of their homecage and kept in a clean holding cage in the behavior testing room, a 3-min aggression screening was started after a male C57BL/6J mouse (screening mouse; 10 weeks) was brought into the homecage of the aggressor, followed by covering the cage with a transparent acrylic lid. During screening, the aggressor freely interacted with the screening mouse. The aggressor was brought back to the holding room after the screening mouse was taken away from the aggressor’s homecage and the female and pups were brought back to its homecage right after screening. Aggressors were selected for trauma induction on Day0 if they showed biting attacks on all of these screening days and the latencies to the initial bites on Day-3 and Day-1 were less than 20 s.

### Trauma induction (Day0)

The following experimental assay emotionally introduced an acute traumatic experience in mice through a social process. The setup was the aggressor’s homecage, divided into an 80×180-mm auditorium zone and a 300×180-mm battle arena by the insertion of a stainless-steel mash with 8×8-mm lattices. The cage was covered with a transparent acrylic lid. The behavioral procedure was carried out in the behavior testing room during CT 4:00 – 6:00, with 3 – 5 experiments done in parallel.

After the female and pups with the aggressor were taken out of their homecage, a divider was inserted into the aggressor’s homecage, allowing the aggressor to freely behave in the battle arena, but not to enter the auditorium zone. A 5-min aggression encounter session started after the focal mouse was brought to the auditorium zone and its partner to the battle arena. Tail rattling counts of the focal mouse during aggressive encounter were recorded by experimenter. The aggressive encounter session was followed by a 5-min stress infiltration session, in which the partner was brought to the focal mouse in the auditorium zone, while the aggressor remained in the battle arena. Right after the stress infiltration session, both focal mouse and its partner were brought back to their homecage in the behavior testing room for a 10-min resting period. The procedure was repeated 5 times with different aggressors. During each resting session, the aggressor stayed in its homecage without the divider before its next intruder encounter. Each aggressor had 3 – 5 encounters with resting periods of 10 – 30 min. After the 5^th^ aggression encounter session, the focal mouse was placed back in its homecage where the nest had been razed, and brought back to the holding room. Partners from different pairs were brought to a new holding cage and housed in groups of 3 – 5 per cage. Right after the last intruder encounter for each aggressor, the female and pups were brought back to the homecage and returned to the holding room together with the aggressor.

### Post-traumatic period (Day0 to Day28)

To investigate the behavior of focal mice after trauma induction (now called ParObsIso mice, Figure 2A), they were housed individually for 4 weeks after the procedure (Day0 to Day28). No environmental enrichment was provided, except to the ParObsIsoEE mice, and the holding cage was not changed during social isolation.

### Control experiments

To differentiate behavioral consequences of the emotionally traumatic experience from consequences of social isolation, a control group of mice had their partners taken away and their nests razed during body weighing on Day0 without trauma induction (xScenIso mice, Figure 1B).

To examine the potential reversal effects of social support on the emotionally traumatic experience, a control group of mice was kept pair-housed with their attacked partners after trauma induction (ParObsParPH mice, Figure 1C).

To characterize potential reversal effects through environmental factors besides social factors, a control group of mice was housed individually with environmental enrichment, provided with a pair of InnoDome™ and InnoWheel™ (Bio-Serv, Inc., Flemington, NJ, USA) and a Gummy Bone (Petite, Green; Bio-Serv, Inc.), after trauma induction (ParObsIsoEE mice, Figure 1D).

To demonstrate predictive validity of potential treatment on stress by an antidepressant, a control group of mice was intraperitoneally injected with fluoxetine (2 µl/g of 10 mg/ml fluoxetine hydrochloride dissolved in saline, i.e. 20 mg/kg; F132-50MG; Sigma-Aldrich, Inc., Saint Louis, MO, USA) once per day at CT 1:00 – 2:00 after trauma induction (ParObsIsoFLX mice, Figure 1E).

To further test the critical component of social relationship in the potential social support reversal, a control group of mice was kept pair-housed but with a stranger mouse after trauma induction (ParObsStrPH mice, Figure 1F).

To identify the impacts of aggression during trauma induction, a control group of mice experienced exposure to strangers of the same strain, gender, and age, instead of the aggressor mice for trauma induction (xAggrExpIso mice, Figure 1G).

To test the influence of social relationship on the emotionally traumatic experience, a control group of mice witnessed the traumatic events toward stranger mice of the same strain, gender, and age instead (StrObsIso mice, Figure 1H). In each iteration of the aggression encounter, stress infiltration, and resting period, a different stranger mouse was presented.

To identify anxiety-like spontaneous behaviors putatively induced by somatic uncertainty, a group of mice was initially sedated with 3%v/v isoflurane in oxygen and then intraperitoneally injected with caffeine (20 µl/g of 0.75 mg/ml anhydrous caffeine dissolved in saline, i.e. 15 mg/kg; 06712-55; Nacalai Tesque, Inc., Kyoto, Japan). Recording of spontaneous behaviors were started 30 min after the injections.

To identify anxiety-like spontaneous behaviors putatively induced by cognitive uncertainty, a group of mice was initially sedated with 3%v/v isoflurane in oxygen and then intraperitoneally injected with 2 µl/g saline. The mice received a series of foot shocks (1 mA for 1 s, 6 times in 5 min, i.e. once every 50 s for the first started at 49 s after placed in the chamber; single chamber system; O’Hara & Co., Ltd., Tokyo, Japan) 25 min after the injections. Recording of spontaneous behaviors were started 30 min after the injections.

To identify non-treated-like spontaneous behaviors, a control group of mice was initially sedated with 3%v/v isoflurane in oxygen and then intraperitoneally injected with 2 µl/g saline. Recording of spontaneous behaviors were started 30 min after the injections.

### Body mass and nest wall height

In the holding room, body masses of all individuals were recorded on Days -7, 0, 1, 7, 28, while the heights of nest walls built by each individual were recorded on Days 1, 7, 28. The height of the nest wall was measured with 5-mm resolution using a transparent acrylic ruler, while the mouse was weighed with 10-mg resolution on a balance. Mice were placed back in their homecages right after recording.

### Light-dark box test

The light-dark box test is an experimental assay to measure anxiety in rodents (Crawley and Goodwin, 1980), designed to evaluate their natural aversion to brightly lit areas against their temptation to explore. The light-dark box setup consisted of two connected 200×200×250-mm non-transparent PVC boxes, separated by a wall with a 50×30-mm door. The boxes were covered with lids with white and infrared LED light illumination for the light and dark areas, respectively, and CCD cameras in the centers (4-chamber system; O’Hara & Co., Ltd., Tokyo, Japan). The floors of the boxes were white, while the walls of the boxes were white for the light area and black for the dark area. Uniform illumination in the light area was 550 lux. Behavioral tests were carried out on Days -7, 1, 7, and 28 in the behavior testing room at 22.7 – 23.0 °C, 51 – 54% humidity, -11 – -9 Pa differential pressure, and 53.6 dB(C) ambient noise level during dark period (CT 6:00 – 8:00).

After habituation for 10 min individually in the homecage in the behavior testing room in darkness, the focal mouse was transferred to the dark area through a 50×50-mm side door. A 5-min behavior record was started right after the side door of dark area was closed and the door between light and dark areas was opened. Locomotion was recorded 2-dimensionally at 15 Hz from top-view with CCD video cameras. Right after recording, the mouse was returned to its homecage, and brought back to the holding room.

### Elevated plus-maze test

The elevated plus-maze test is an experimental assay to measure anxiety in rodents (Pellow et al., 1985), designed to evaluate their natural fear of falling and exposure against their temptation to explore. The elevated plus-maze setup consisted of a gray PVC platform raised 500 mm above the ground (single maze system; O’Hara & Co., Ltd.). The platform was composed of a 50×50-mm square central platform, two opposing 50×250-mm open arms, and two opposing 50×250-mm closed arms with 150-mm semi-transparent walls. Each of the two open arms emanated at 90° to each of the two closed arms, and vice versa. The apparatus was installed in a soundproof box with white fluorescent lamp illumination (20 lux) and ventilators. Behavioral tests were carried out on Days -7, 1, 7, 28 in the behavior testing room at 22.8 – 23.0 °C, 53 – 56% humidity, -13 – -11 Pa differential pressure, and 52.1 dB(C) ambient noise level during dark period (CT 8:00 – 10:00).

After habituation for 10 min individually in the homecage in the behavior testing room in darkness, the focal mouse was brought to the central platform of the elevated plus-maze, facing the open arm on the opposite side from the door of the soundproof box. A 5-min behavior recording was started right after the door of the soundproof box was closed. Locomotion was recorded 2-dimensionally at 15 Hz from top-view with a CCD video camera installed above the center of the central platform. Delineated entrances to open and closed arms were defined at 50 mm from the center of the central platform. Right after recording, the mouse was placed back in its homecage, and brought back to the holding room.

### Open field test

The open field test is an experimental assay to measure anxiety in rodents (Hall and Ballachey, 1932), designed to evaluate their spontaneous activity under a gradient of spatial uncertainty (high in the field center and low along the walls and at the corners of the field). The open field setup consisted of a 400×400×300-mm non-transparent gray PVC box with no cover, installed in a soundproof box with white LED light illumination and ventilators (2-chamber system; O’Hara & Co., Ltd.). Behavioral tests were carried out on Days -7, 1, 7, 28 in the behavior testing room at 22.8 – 23.0 °C, 53 – 56% humidity, -13 – -11 Pa differential pressure, and 56.7 dB(C) ambient noise level during dark period (CT 8:00 – 10:00).

After habituation for 10 min individually in the homecage in the behavior testing room in darkness, the focal mouse was brought to the center of the open field arena under 20-lux uniform illumination, facing the wall on the opposite side from the door of the soundproof box. A 5-min behavior recording was started right after the door of the soundproof box was closed. Locomotion was recorded 2-dimensionally at 15 Hz from top-view with a CCD video camera installed above the center of the open field arena. Vertical activity of exploratory rearing behavior was recorded by the blocking of invisible infrared beams created and detected by photocell emitters and receptors, respectively, positioned 60 mm high on the walls of the open field box. A delineated center region was defined as the central 220×220 mm area. Right after recording, the mouse was placed back in its homecage, and returned to the holding room.

### Locomotor activity test

The locomotor activity test is an experimental assay to measure spontaneous activity of rodents in an environment without an experimentally designed stressor. The locomotor activity setup consisted of a 200×200×250 mm non-transparent covered PVC box with infrared LED illumination and a CCD camera in the center (the dark area of the light-dark box setup, while the door between the light and dark areas was closed and fixed). The floor of the box was embedded with bedding material from the homecage of the focal mouse, while the walls of the box were black. Behavioral test was carried out on Days -7, 1, 7, 28 in the behavior testing room at 22.7 – 23.0 °C, 51 – 54% humidity, -11 – -9 Pa differential pressure, and 53.6 dB(C) ambient noise level during dark period (CT 6:00 – 8:00).

After habituation for 30 min individually in the behavior testing box, a 1-hr behavior recording was started. The behavior testing box was not covered completely in order to allow air circulation. Locomotion was recorded 2-dimensionally at 15 Hz from top-view with the CCD video camera. Right after recording, the mouse was returned to its homecage, and brought back to the holding room.

### Active social contact test

The active social contact test [also known as “social interaction test”, but to be distinguished with the one-session test using an open field with a social target freely behaving in the field (Arakawa et al., 2014) or the one-session test placing a social target-containing cylinder into the center of the testing subject’s homecage for social instigation (Tsuda and Ogawa, 2012)] is a 2-session experimental assay to measure social motivation in rodents (Berton et al., 2006). The setup consists of a 400×400×300-mm non-transparent gray PVC box with no cover, installed in a soundproof box with 20-lux white LED illumination and ventilators.

A 60×100×300-mm stainless-steel chamber with wire grid sides was placed in the center of the wall on the opposite side from the door of the soundproof box. The wire grid had 8×8 mm lattices at a height of 10 – 60 mm from the bottom. An ultrasound microphone (CM16/CMPA; Avisoft Bioacoustics, Glienicke, Germany) with an acoustic recording system (UltraSoundGate; Avisoft Bioacoustics) was hung outside the chamber, 100 mm above the ground. Behavioral tests were carried out on Days -7, 1, 7, 28 in the behavior testing room at 22.8 – 23.0 °C, 53 – 56% humidity, -13 – -11 Pa differential pressure, and 56.7 dB(C) ambient noise level during dark period (CT 8:00 – 10:00).

The social target used for active social contact tests was either a male or a female C57BL/6J mouse (18 weeks), pair-housed with a partner of the same strain, gender, and age for more than 2 weeks before the tests. The social target was adapted to the experimental protocol one day before the tests in the behavior testing room during dark period (CT 8:00 – 9:00): After habituation for 5 min individually in the homecage in the soundproof box under 20-lux uniform illumination, the social target was brought into the chamber in the open field arena under 20 lux uniform illumination. A male C57BL/6J mouse (11–16 weeks; from partners of xScenIso mice in previous experiment) was then brought to the open field arena for a 2.5-min spontaneous exploration and interaction with the social target. The social target was then brought back to its homecage in the soundproof box under 20-lux uniform light for a 5-min rest. The social interaction procedure was repeated with a different male C57BL/6J mouse right afterward. After the social target had interacted with 4 different mice, it was returned to its homecage and brought back to the holding room.

On testing days, after 10-min habituation individually in its homecage in the behavior testing room in darkness, the first session of the active social contact test started by placing the focal mouse at the center of the open field arena under 20-lux uniform light, facing the empty chamber. A 2.5-min behavior recording started right after the door of the soundproof box was closed. Locomotion was recorded 2-dimensionally at 15 Hz from top-view with a CCD video camera installed above the center of the open field arena. Ultrasonic vocalization was recorded at 250 kHz. In the second session of the active social contact test, which followed the first session, the social target was brought into the chamber. Another 2.5-min behavior recording started as soon as the door of the soundproof box was closed. Right afterward, the focal mouse was returned to its homecage and brought back to the holding room.

The focal mouse experienced active social contact tests with different social targets on different recording days (Days -7, 1, 7, 28), while different focal mice were tested with the same social target on the same recording day (5 – 10 records). The social target remained in its homecage in a soundproof box under 20-lux uniform illumination before and between each test. A delineated interaction zone was taken as the region within 80 mm of the edges of the chamber. Social approaches of the focal mouse poking its nose toward the social target were recorded manually using the event recording software, tanaMove ver0.09 (http://www.mgrl-lab.jp/tanaMove.html).

### Partner-revisiting test

The partner-revisiting test is a memory-based experimental assay to measure social bonding in rodents [sharing similar concept of “familiar *v.s.* novel social target recognition”, but to be distinguished with the three-chamber paradigm test (Nadler et al., 2004)]. The partner-revisiting setup was the uncovered homecage of the focal mouse, installed in a soundproof box with white LED illumination and ventilators (O’Hara & Co., Ltd., Tokyo, Japan). The long sides of the homecage were parallel to the door of soundproof box. The partner-revisiting test was carried out on Day28 in the behavior testing room at 22.8 – 23.0 °C, 53 – 56% humidity, -13 – -11 Pa differential pressure, and 56.7 dB(C) ambient noise level during light period (CT 4:00 – 6:00) with 350-lux light intensity.

The previously separated partner of the focal mouse, being a social target in the test, was initially sedated with 3%v/v isoflurane in oxygen, and then anesthetized by intraperitoneal (i.p.) injection of a mixture of medetomidine (domitor, 3%v/v in saline, 0.3 mg/kg), midazolam (dormicum, 8%v/v in saline, 4 mg/kg), and butorphanol (vetorphale, 10%v/v in saline,5 mg/kg). Also, a stranger mouse (15 weeks; a separated partner of a ParObsIso or Buffered mouse for testing a xScenIso, StrObsIso, or xAggrExpIso mouse, and *vice versa*) was anesthetized as an alternative social target. Both anesthetized mice were kept on a heating pad at 34°C (B00O5X4LQ2; GEX Co., Ltd., Osaka, Japan) to maintain their body temperatures before the test.

The focal mouse was brought to a clean, uncovered holding cage in the soundproof box under 50-lux uniform illumination for 5-min habituation, while its homecage was placed in another soundproof box under 50-lux uniform light. During habituation of the focal mouse, the anesthetized social targets were injected with atipamezole hydrochloride (antisedan; 6%v/v in saline for 0.3 mg/kg, i.p.) to induce recovery from anesthesia. During the waking-up period, the social targets were still immobilized and not able to actively interact with the focal mouse during the following recording, but showed enough social cues to be attractive for the focal mouse. The immobilized social targets were then placed in the homecage of the focal mouse with their nose pointing toward the center of the short side of the wall (10 mm of nose-to-wall distance) with their bellies facing the door of the soundproof box. After habituation, the focal mouse was brought to the center of its homecage, facing the long side of the homecage wall on the opposite side from the door of soundproof box. A 10-min behavior record started right after the door of the soundproof box was closed. Locomotion was recorded 2-dimensionally at 15 Hz from top-view with a CCD video camera installed above the center of the homecage. Right after recording, social targets were taken out of the focal mouse’s homecage and the focal mouse was brought back to the holding room.

Social contacts including sniffing, allogrooming, and pushing of the focal mouse toward each of the social targets were recorded manually using the event recording software, tanaMove ver0.09 (http://www.mgrl-lab.jp/tanaMove.html).

### Baseline plasma corticosterone concentration test

The baseline plasma corticosterone (CORT) concentration test is a competitive-inhibition enzyme-linked immunosorbent assay (ELISA) to measure physiological stress level in rodents, designed to quantitatively determinate CORT concentrations in blood plasma. The sample collection was carried out on Days -7, 1, 7, 28 in the behavior testing room at 22.4 – 23.0 °C, 53 – 58% humidity, -4 – -3 Pa differential pressure, and 57.1 dB(C) ambient noise level during CT 4:00 – 6:00 with 350-lux white light.

After habituation for 30 min individually in the homecage in the behavior testing room, the mouse was initially sedated with 3%v/v isoflurane in oxygen. Six drops of blood from the facial vein pricked by a 18G needle were collected in a EDTA-lined tube [K2 EDTA (K2E) Plus Blood Collection Tubes, BD Vacutainer; Becton, Dickinson and Company (BD), Franklin Lakes, NJ, USA] and kept on ice. Right after collection, the mouse was returned to its homecage, and brought back to the holding room. Whole blood samples were then centrifuged (MX-300; Tomy Seiko Co., Ltd., Tokyo, Japan) at 3,000 rpm for 15 min at 4°C. Plasma supernatant was decanted and kept at -80°C until the measurement on Day 29.

CORT concentrations in blood plasma were tested with Mouse Corticosterone (CORT) ELISA Kit (MBS703441, 96-Strip-Wells; MyBioSource, Inc., San Diego, USA; stored at 4°C before use) on Day 29. All reagents [assay plate (96 wells, pre-coated with goat-anti-rabbit antibody), standards (0, 0.1, 0.4, 1.6, 5, and 20 ng/ml of CORT), rabbit-anti-CORT antibody, HRP-conjugated CORT, concentrated wash buffer (20x phosphate-buffered saline (PBS)), 3,3’,5,5’-tetramethylbenzidine (TMB) color developing agent (substrates A and B), and TMB stop solution] and samples were brought to room temperature for 30 min before use. Collected plasma samples after thawing were centrifuged again (MX-300; Tomy Seiko Co.) at 3,000 rpm for 15 min at 4°C. 20 μl of standard or sample was added per well, assayed in duplicate, with blank wells set without any solution. After 20 μl of HRP-conjugated CORT was added to each well except to the blank wells, 20 μl of rabbit-anti-CORT antibody was added to each well and mixed. After incubation for 1 hour at 37°C, each well was aspirated and washed, repeated for 3 times, by filling each well with 200 μl of wash buffer (diluted to 1x PBS) using a squirt bottle, standing for 10 s, and completely removing liquid at each step. After the last wash and the removal of any remaining wash buffer by decanting, the plate was inverted and blotted against clean paper towels. After TMB color developing agent (20 μl of substrate A and 20 μl of substrate B) was added to each well, mixed, and incubated for 15 min at 37°C in dark, 20μl of TMB stop solution was added to each well and mixed by gently tapping the plate. The optical density (O.D.) of each well was determined, within 10 min, using a microplate reader (Multiskan GO; Thermo Fisher Scientific, Inc., Waltham, MA, USA) measuring absorbance at 450 nm, with correction wavelength set at 600–630 nm.

CORT concentrations were calculated from the O.D. results using custom scripts written in MATLAB R2015b (MathWorks). The duplicate O.D. readings for each standard and sample was averaged and subtracted the average O.D. of the blanks, *X* = 〈*O*. *D*.〉 − 〈*O*. *D*.〉_*blank*_. A standard curve was determined by a four parameter logistic (4PL) regression fitting the equation 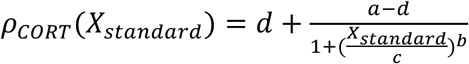, where *ρ_CORT_* is the CORT concentration, *a* is the minimum asymptote, *b* is the Hill’s slope, *c* is the inflection point, and *d* is the maximum asymptote. CORT concentrations of the samples were calculated from the fitted 4PL equation with respected to *X_sample_*.

### Ex vivo diffusion tensor imaging

*Ex vivo* diffusion tensor imaging (DTI) is a magnetic resonance imaging (MRI) technique to determinate structural information about tissues (Basser et al., 1994), designed to measure the restricted diffusion of water in tissue. The sample collection was carried out on Day 28 in the necropsy room at 22.4 - 22.5 °C, 53 - 54 % humidity, and 10 - 12 Pa differential pressure during CT 4:00 - 6:00 with 750-lux white light.

After mice were brought individually in their homecages to the necropsy room, they were initially sedated with 3%v/v isoflurane in oxygen, then deeply anaesthetized with a ketamine-xylazine mixture (>30 μl/g body weight of 100 mg/ml ketamine and 20 mg/ml xylazine), and perfused transcardially. The perfusates, in a two-step procedure, were (i) 20 ml of ice cold 1x phosphate-buffered saline (PBS), and (ii) 20 ml of ice cold 4% paraformaldehyde, 0.2% sodium meta-periodate, and 1.4% lysine in PBS. Mouse skull including the brain was removed and stored in the perfusate (ii) at 4 °C for 2 weeks. Each skull with the brain was then transferred into 2 mM gadolinium with diethylenetriaminepentaacetic acid (Gd-DTPA) and 0.5% azide in PBS for 2 weeks.

Isolated fixed brains within the skulls were positioned in an acrylic tube filled with fluorinert (Sumitomo 3M Ltd., Tokyo, Japan) to minimize the signal intensity attributable to the medium surrounding the brain during MRI scanning. All MRI was performed with an 11.7-T MRI system (BioSpec 117/11; Bruker Biospec, Ettlingen, Germany) using ParaVision 6.0.1 software (Bruker Biospec, Ettlingen, Germany) for data acquisition. The inner diameter of the integrated transmitting and receiving coil (Bruker Biospec, Ettlingen, Germany) was 35 mm for the ex vivo MRI. DTI data were acquired by using a 3-D diffusion-weighted spin-echo imaging sequence, with repetition time (TR) = 267 ms, echo time (TE) = 18.5 ms, b-value = 2,000 s/mm2, and 30 non-collinear directions. Five T2-weighted measurements were acquired together with DTI, for one every 6 diffusion measurements. The acquisition matrix was 216×216×168 over a 27.0×27.0×21.0 mm3 field of view, resulting in a native isotropic image resolution of 125 μm. Total acquisition time was 96 hr.

MRI data was processed using custom scripts written in MATLAB R2015b (MathWorks). All 30 DTI and 5 T2 3-D images were masked by thresholding at the half of mean values of diffusion weights for each voxel and omiting clusters smaller than 10 voxels. After diffusion tensor of each voxel was estimated by solving the Stejskal-Tanner equation through linear regression (Hrabe et al., 2007), the 3 eigenvalues (*λ*_1_, *λ*_2_, and *λ*_3_) with respect to the 3 axes of the diffusion ellipsoid (the longest, middle, and shortest axes, respectively) were calculated by eigenvalue decomposition of the diffusion tensor. Four focused DTI-based measures (Mori, 2007) are the mean diffusivity (MD) that represents membrane density

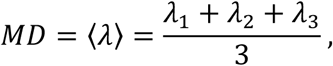

axial diffusivity (AD) that represents neurite organization

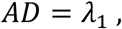

radial diffusivity (RD) that represents myelination

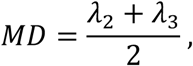

and fractional anisotropy (FA) that represents average microstructural integrity

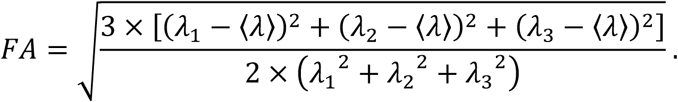

After the registration of these 3-D DTI-based brain maps to a template brain atlas (DSURQE Atlas; https://wiki.mouseimaging.ca/display/MICePub/Mouse+Brain+Atlases), mean values of these DTI-based quantities were identified in a total of 244 brain regions ($) for each individual. Based on the FA maps, DTI-based long-range fiber tracking from a focal seed region ($B) was calculated with 2 samplings, distance of forward fiber in one step = 50 μm, and thresholds of minimal fiber length = 750 μm, maximal fiber length = 75,000 μm, maximal fiber deviation angle = 57.3°, and minimal FA for keeping tracking = 0.4.

### Video data processing

Video image data was processed using custom scripts written in MATLAB R2015b (MathWorks). Each video frame from a recorded AVI video file was read as a 2-dimensional matrix with an 8-bit gray scale. Each of these matrices was then divided by a background matrix read from a TIF image file of the background taken before bringing the test mouse to the setup. The centroid of the area with non-one values in each matrix ratio was taken as the position of the mouse at this specific time point. Speed was calculated as the distance between temporally adjacent positions multiplied by 15 (15-Hz recording). Freezing periods were sorted out if the area of the mouse body between temporally adjacent frames was less than 20 mm^2^.

### Audio data processing

Audio signal data was processed with custom scripts written in MATLAB R2015b (MathWorks). Each recorded WAV audio file was read and transformed into a spectrogram using fast Fourier transform with non-overlapping 0.4-ms time windows. To identify the time segments with ultrasonic vocalization signals, recordings were thresholded at a power spectral density (PSD) ≥-75 dB/Hz, and time segments with averaged PSD between 0–50 kHz higher than that between 50–120 kHz were removed. The duration of remaining time segments was calculated.

### Statistical analysis

Numerical data were analyzed with custom scripts written in MATLAB R2015b (MathWorks). Statistical significance of the difference between 2 mean values was estimated with two-tailed, two-sample Student’s t-test, except of DTI quantities and state-space behavioral comparison which were estimated with one-tailed, two-sample Student’s t-test. Statistical significance of the difference between 2 median values (vocalization analysis; Figure 8F) was estimated using one-tailed Mann–Whitney U test.

To capture fine-scale behavioral details of location within the light-dark box and the elevated plus-maze (Figures 4 and 5), we computed T(x), the cumulative probability of finding position ≤x, for each individual (light traces) for all measured locations (a collection of locations from all mice for the statistics). We then show the average across the control group (bold blue trace) and the ParObsIso group (bold red trace). We compared the averages of each group with a two-tailed, two-sample Student’s t-test and plot the resulting p-values, presented as -log(p), the negative logarithm of p-values. We also show the box plot (the minimum, lower quartile, median, upper quartile, and maximum) of -log(p) values collapsed across all measured locations. To capture the fine-scale behavioral details of speed, we followed a similar procedure as above, but with U(v), the cumulative distribution function of finding speed ≤v.

To estimate local likelihoods of caffeine-injected, foot-shocked, and non-treated behavior in the light-dark box or elevated plus-maze tests for any given 4-dimensional behavioral states described by the position, speed, velocity along the stressor axis, and acceleration strength, we trained a deterministic 3-layer feedforward network with hidden layer sizes of 26, 30, and 24 units, respectively, using log-sigmoid transfer functions. For pattern recognition, each network was trained by using the scaled conjugate gradient method to minimize cross-entropy to obtain reliable classifiers, with a random data division of 80% for training and 20% for testing. Training of updating weights and biases terminated when one of the following condition was matched: (1) reaching 1,000 iterations, (2) obtaining a perfect data fitting [i.e. the mean squared error (MSE) equaled to zero], (3) having the error rate continuously increasing for more than 6 epochs, (4) showing the gradient of MSE less than 10^-7^, and (5) receiving the training gain larger than 10^10^. The global likelihoods of a recorded mouse to be caffeine-injected-like, foot-shocked-like, and non-treated-like were calculated by taking the average of local likelihoods of each experimental type estimated by the corresponding trained network.

To evaluate the uncertainty of the percentage for each tail rattle count (Figure 10A), we created 10,000 bootstrapped data sets where each sample was randomly picked with replacement from the original data set. Each bootstrapped data set had the same sample size as the original data set. The standard error was taken as the standard deviation of the bootstrapped percentages for a tail rattle count. A similar procedure was carried out to evaluate the standard error of mean for the percentage of time spent in each behavior in the partner-revisiting test (Figures 10H and 11E), where each sample of the bootstrapped data sets was a set of the percentages of the three classified behaviors (partner concern, stranger concern, and non-social activity/sniffing at social targets) from a mouse record. Standard errors of means for other results were estimated with the formula 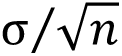, where σ is the sample standard deviation and *n* is the sample size.

## Acknowledgments

We thank Tsuyoshi Koide (NIG, Japan), Aki Takahashi (University of Tsukuba), Teruhiro Okuyama (MIT), Charles Yokoyama (RIKEN BSI), Ai Koizumi (CiNet), and Tsai-Wen Chen (National Yang-Ming University) for comments, Tadashi Yamamoto (OIST) for sharing behavior testing setups, and Steven D. Aird (OIST) for editing the manuscript.

## Author contributions

R.X.L. conceptualized the study, designed the research, performed experiments, analyzed data, interpreted results, wrote the original draft, and revised the article; G.J.S. advised on data analysis and revised the article; B.K. advised on research design and revised the article.

---

**Supplemental Figure 1.**
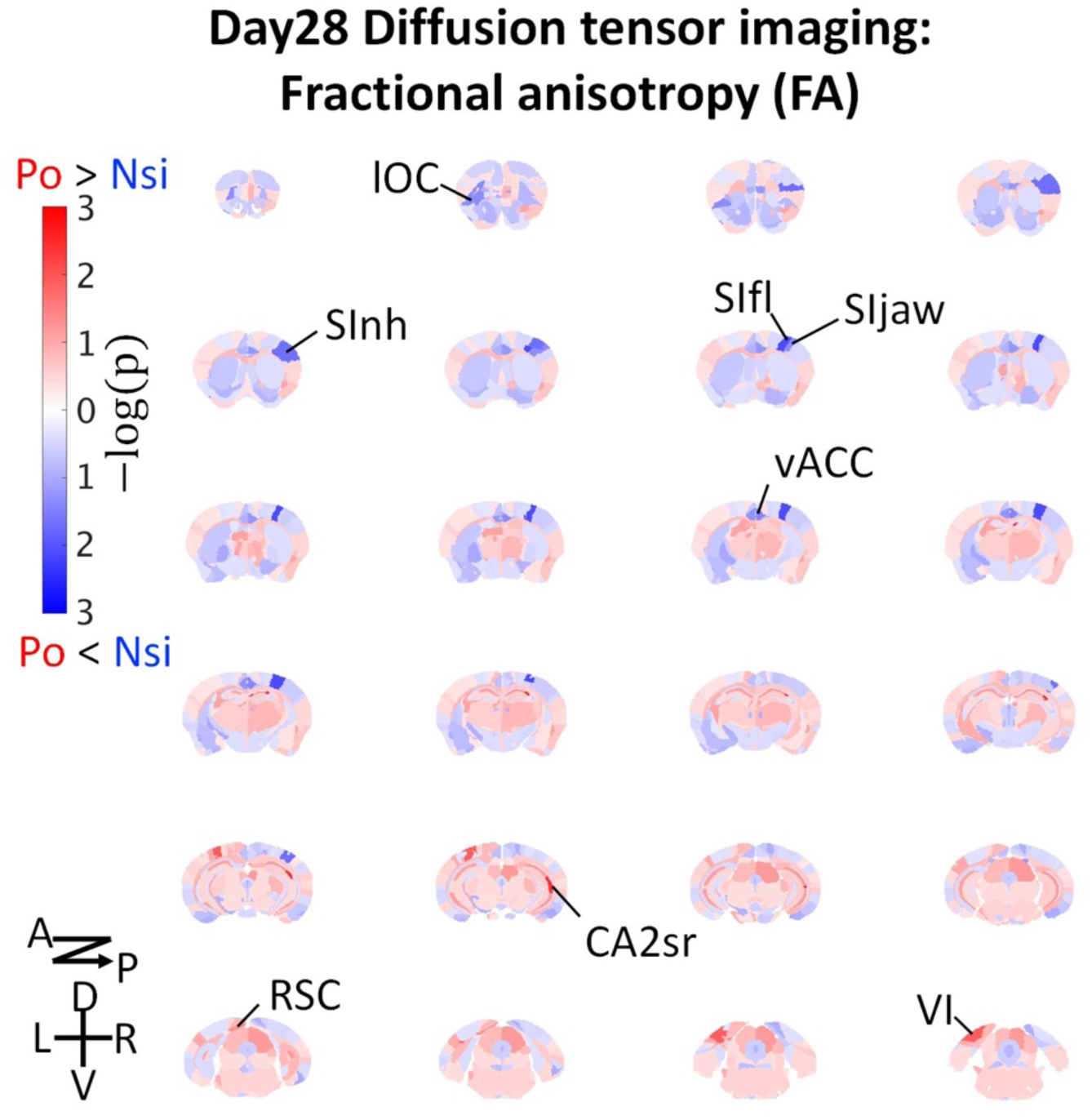
Brain-wide microstructural changes measured by DTI fractional anisotropy. Po, ParObsIso mice; Nsi, xScenIso mice; -log(p), statistical significance through a two-population Student’s t-test; A, anterior; P, posterior; D, dorsal; V, ventral; L, left; R, right; lOC, lateral orbital cortex; SInh, non-homunculus region of the primary sensory cortex; SIfl, forelimb region of the primary sensory cortex; SIjaw, jaw region of the primary sensory cortex; vACC, ventral region of the anterior cingulate cortex; CA2sr, stratum radiatum of the hippocampal cornu ammonis (CA) 2 area; RSC, the retrosplenial cortex; VI, the primary visual cortex.

**Supplemental Figure 2.**
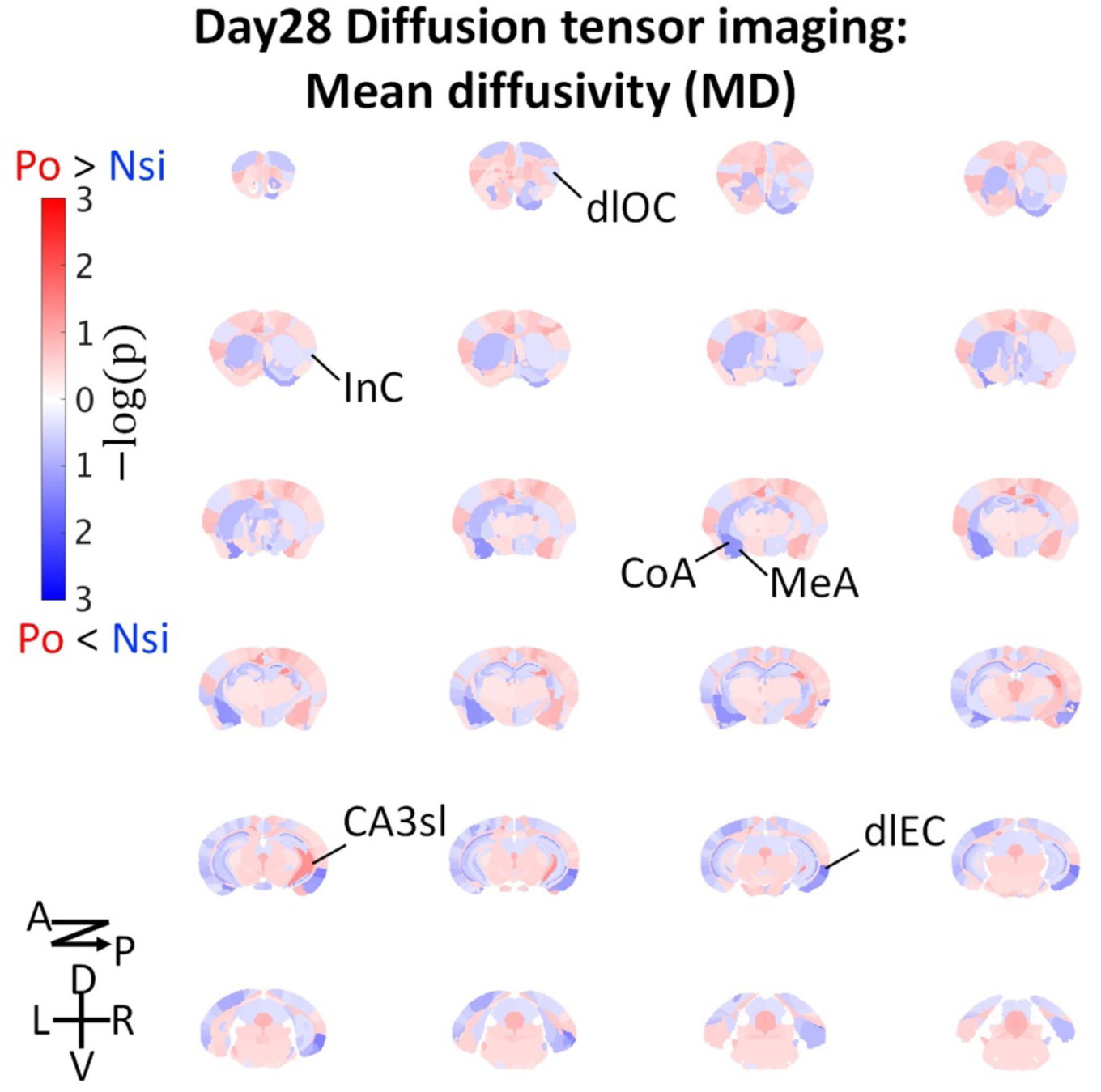
Brain-wide microstructural changes measured by DTI mean diffusivity. dlOC, dorsolateral orbital cortex; InC, the insular cortex; CoA, the cortical amygdalar nucleus; MeA, medial amygdalar nucleus; CA3sl, stratum lucidum of the hippocampal CA3 area; dlEC, dorsolateral entorhinal cortex.

**Supplemental Figure 3.**
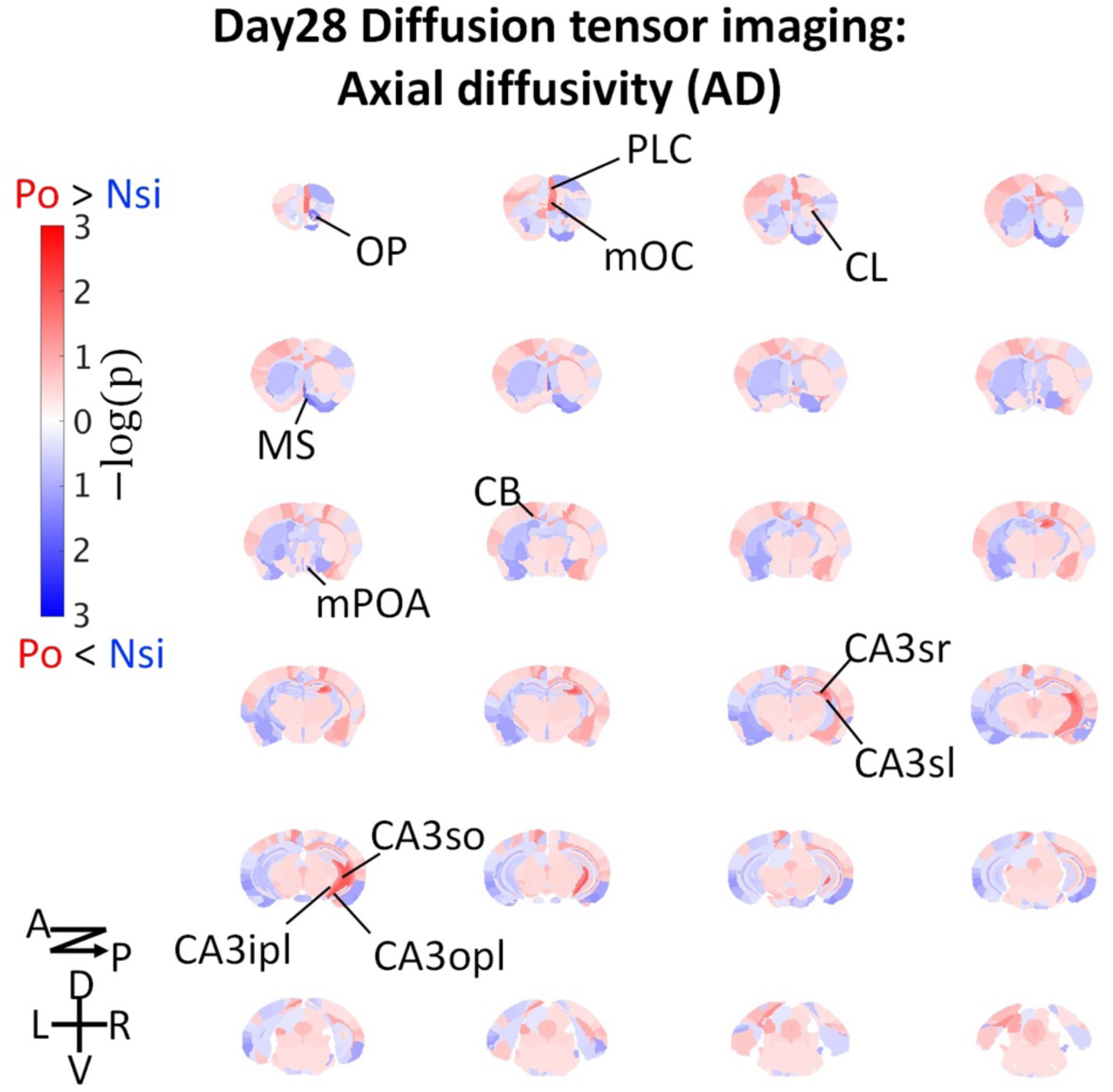
Brain-wide microstructural changes measured by DTI axial diffusivity. OP, olfactory peduncle; PLC, prelimbic cortex; mOC, medial orbital cortex; CL, claustrum; MS, medial septal complex; mPOA, medial preoptic area; CB, cingulum bundle; CA3sr, stratum radiatum of the hippocampal CA3 area; CA3sl, stratum lucidum of the hippocampal CA3 area;CA3so, stratum oriens of the hippocampal CA3 area; CA3ipl, inner pyramidal layer of the hippocampal CA3 area; CA3opl, outer pyramidal layer of the hippocampal CA3 area.

**Supplemental Figure 4.**
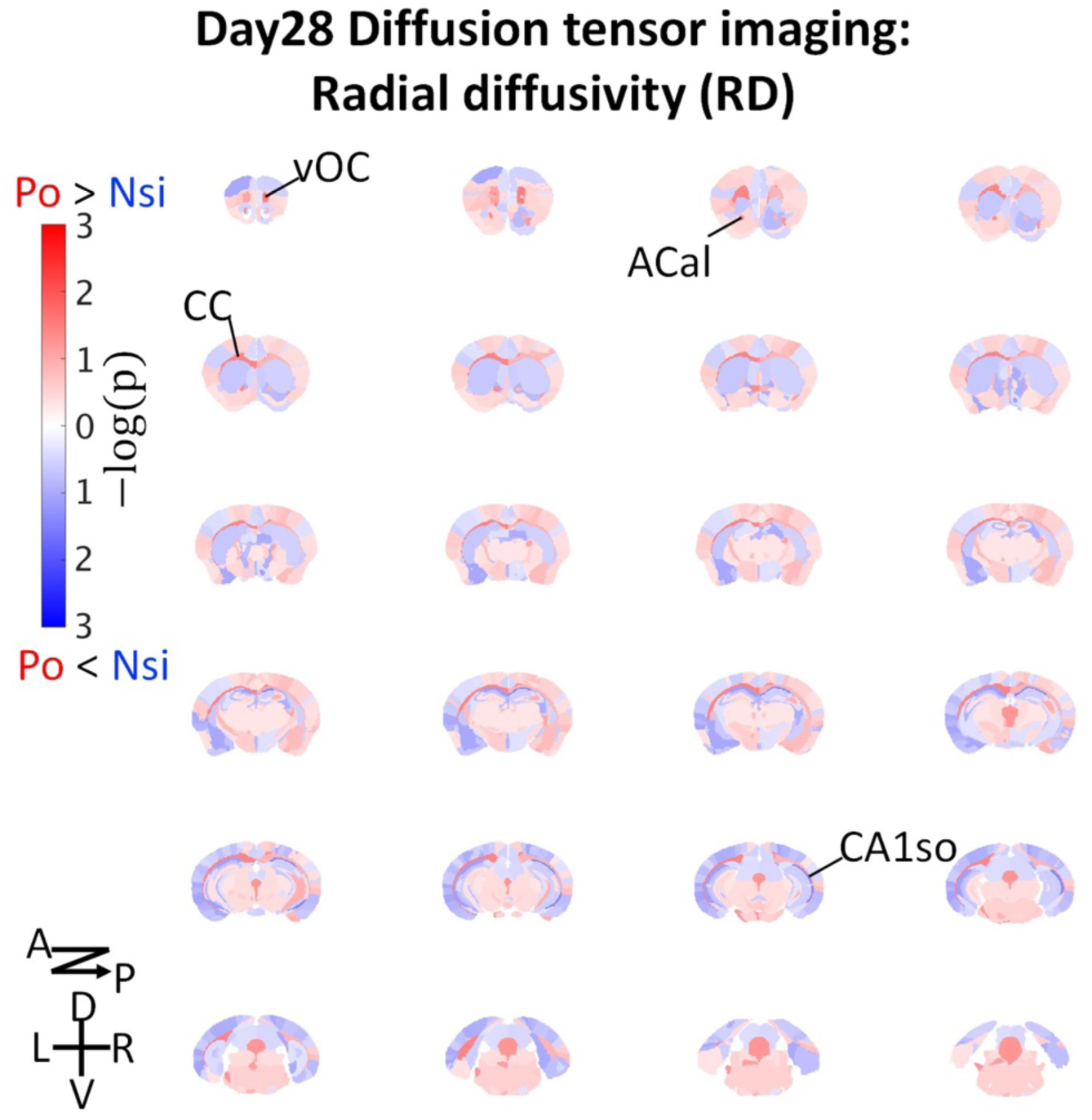
Brain-wide microstructural changes measured by DTI radial diffusivity. vOC, ventral orbital cortex; ACal, anterior limb of the anterior commissure; CC, corpus callosum; CA1so, stratum oriens of the hippocampal CA1 area.

**Supplemental Figure 5.**
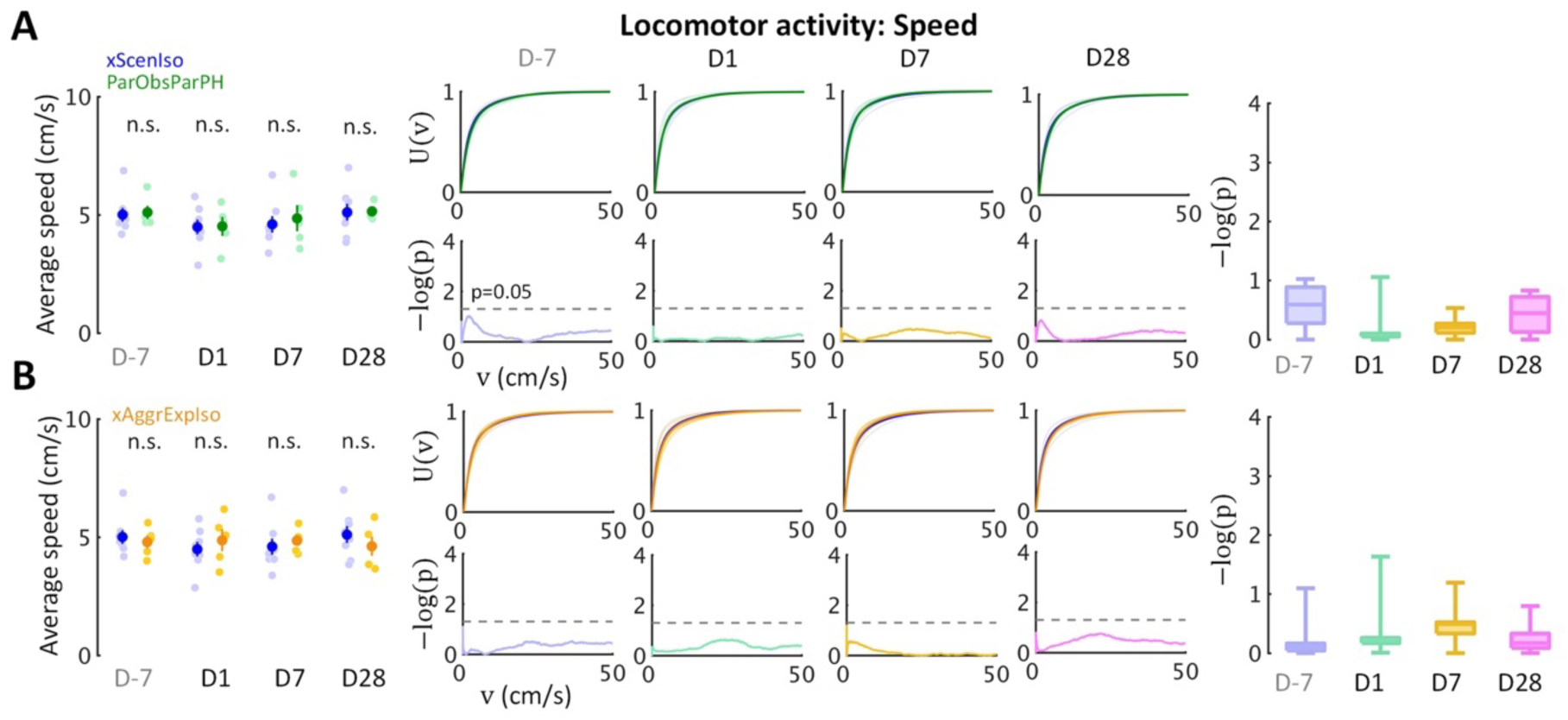
Locomotor activity tests in control experiments agree with the conclusions of spontaneous behaviors given from the light-dark box tests. **(A)** The results of ParObsIsoFLX mice. **(B)** The results of ParObsStrPH mice.

**Supplemental Figure 6.**
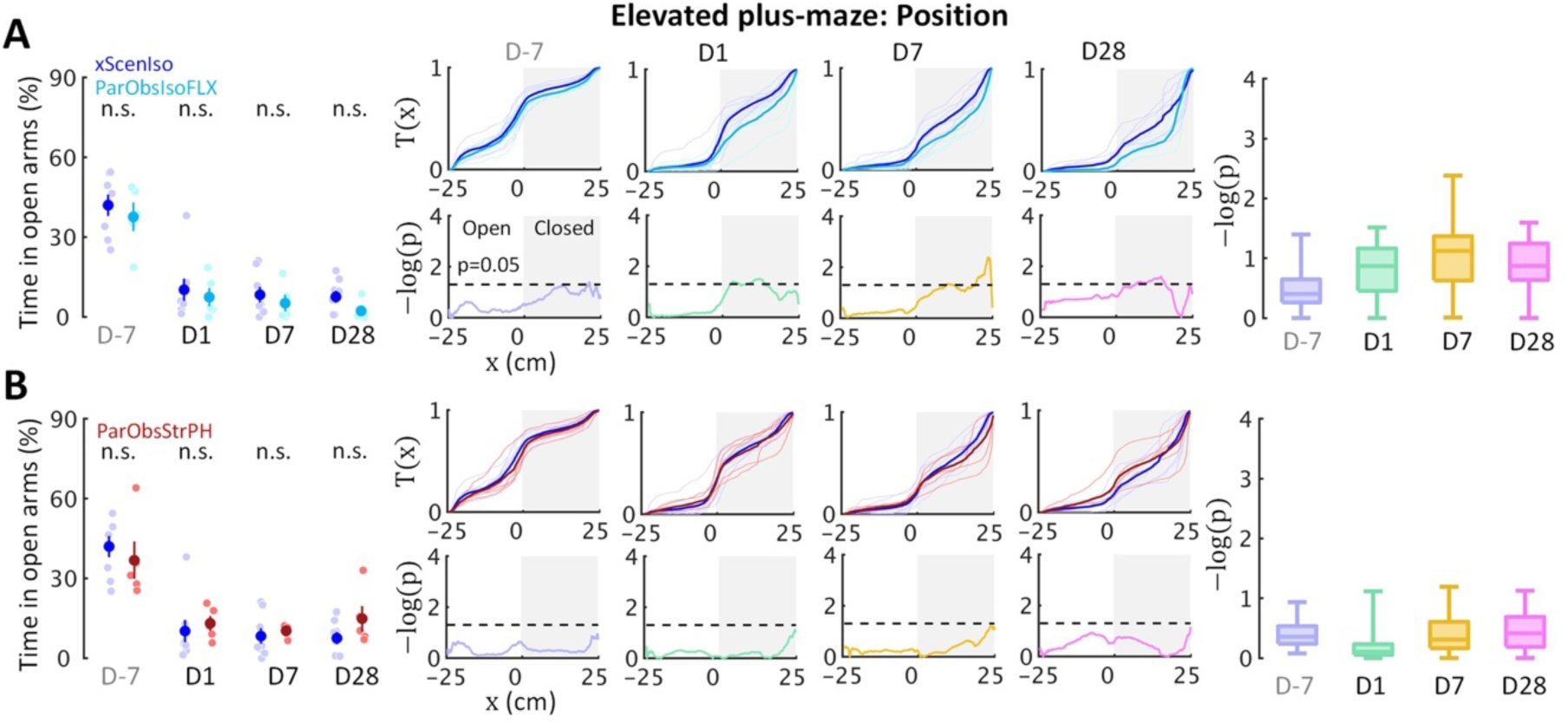
Elevated plus-maze tests in control experiments agree with the conclusions of spontaneous behaviors given from the light-dark box tests. **(A)** The results of ParObsParPH mice. **(B)** The results of xAggrExpIso mice.

**Supplemental Video 1.** Landscape in the 4-D state space of local likelihood of caffeine-injected behaviors in the light-dark box test.

**Supplemental Video 2.** Landscape in the 4-D state space of local likelihood of foot-shocked behaviors in the light-dark box test.

**Supplemental Video 3.** Landscape in the 4-D state space of local likelihood of control behaviors in the light-dark box test.

**Supplemental Video 4.** Landscape in the 4-D state space of local likelihood of caffeine-injected behaviors in the elevated plus-maze test.

**Supplemental Video 5.** Landscape in the 4-D state space of local likelihood of foot-shocked behaviors in the elevated plus-maze test.

**Supplemental Video 6.** Landscape in the 4-D state space of local likelihood of control behaviors in the elevated plus-maze test.

**Supplemental Video 7.** Social apathy is an observed behavioral characteristic of ParObsIso mice. Examples of 30-s recordings during the social session in the female stranger test on Day1. First scene, xScenIso mouse; Second scene, ParObsIso mouse.

**Supplemental Video 8.** Illustration of behavioral characteristics that were specific to observer mice during trauma induction when their partners were attacked. First scene, tail rattling during aggressive encounter; Second scene, tail rattling during aggressive encounter (4x slower); Third scene, hiding under bedding material with the partner during resting. These behaviors were not observed if a stranger mouse got attacked.

**Supplemental Video 9.** Illustration of rebound reaction of a ParObsIso mouse to its previously pair-housed partner during the partner-revisiting test.

